# Discovery and Repurposing of Multi-Target Senolytics through Structure-Based Virtual Screening

**DOI:** 10.1101/2024.07.09.602796

**Authors:** Samael Olascoaga, Mina Königsberg, Francisco Cortés-Benítez, Jaime Pérez-Villanueva, Norma Edith López-Diazguerrero

## Abstract

Cellular Senescence is a state of irreversible cell cycle arrest in response to various stressors that can damage the cell. Senescent Cells (SCs) exhibit multiple alterations at the morphological and molecular levels, one of the most significant being the development and activation of Senescent Cell Anti-Apoptotic Pathways (SCAPs). Due to this characteristic, SCs accumulate in organs and tissues during aging. The accumulation of these cells has been associated with the onset and progression of various chronic degenerative diseases, and their selective elimination allows for the slowing down, halting, and reversing of many age-associated ailments. Small molecules called senolytics, which inhibit SCAPs, have been proposed to selectively eliminate SCs. Herein, we identified new senolytics through computational and rational drug design approaches. Among the identified molecules are the FDA-approved drug tolvaptan, the experimental Phase II drug sotrastaurin, and the experimental drugs cryptotanshinone and bicuculline. The effectiveness of these molecules in targeting senescent cells was confirmed through experiments using two different models of cellular senescence in human lung fibroblasts. Our results suggest that some molecules work by selectively inducing apoptosis through a multi-target mechanism, inhibiting several SCAPs, including PIK3CD, SERPINE1, EFNB1, and PDGFB. These newly identified FDA-approved and experimental drugs have the potential to be repurposed as new senolytic agents.

## Introduction

Cellular Senescence (CS) is an irreversible state of cell cycle arrest in response to various stressing agents that can damage the cell. Senescent cells (SCs) exhibit diverse morphological and molecular alterations, such as the acquisition of an elongated and flattened phenotype in cell cultures, accumulation of lysosomes and misfolded proteins in the cytoplasm, alterations in mitochondrial dynamics, dysfunction in autophagy, among others. In addition to these changes, there are aberrations in gene expression and the secretion of cytokines, chemokines, growth factors, and metalloproteinases into the extracellular medium, collectively known as the senescence-associated secretory phenotype (SASP), which can influence other cells in a paracrine manner and promote a pro-inflammatory environment [1]. An important characteristic of SCs is their ability to resist and evade programmed cell death due to the development and activation of senescent cell anti-apoptotic pathways (SCAPs). This resistance is believed to contribute to accumulating SCs in organs and tissues over the years, promoting the onset and progression of various chronic degenerative diseases [2]. Some signaling pathways, such as PI3K/Akt, MAPK-NF-κB, and Insulin/IGF, as well as certain proteins like FAK, MVP, HSP90, and proteins of the Bcl-2 family, have been identified as components of SCAPs [3].

Several genes have been identified that, when individually inhibited through RNA interference (siRNA), induce selective death of SCs. Some identified genes include ephrins (EFNB1 or 3), PIK3CD, CDKN1A, SERPINE1, PDGFB, BCL-xL, and SERPINEB2. The discovery of these genes involved in SCAPs led to the developing of the first generation of senolytics: Dasatinib (D) and Quercetin (Q). These small molecules are capable of selectively eliminating senescent cells [4].

Unlike the traditional one-drug, one-target strategy, the first senolytics were specifically designed to target multiple SCAPs networks to affect cellular survival. These early senolytics often have diverse pharmacological mechanisms of action, acting synergistically. Clear examples of this approach include the flavonoid fisetin, and the combination of D + Q, both of which have been widely researched. The precise mechanism of action of the D + Q combination is not fully known (as is the case with most senolytics), however, it demostrates potent senolytic activity by interfering with multiple pro-survival signaling networks, including Ephrin-Eph receptor signaling pathways, the PI3K-AKT pathway, and BCL-2 family proteins [5].

The selective elimination of SCs using senolytics could contribute to stopping, slowing, or reversing the onset and progression of various chronic-degenerative diseases associated with aging, such as spinal degeneration [6], renal complications in diabetes [7], arteriosclerosis [8], pulmonary fibrosis [9] and adipose tissue inflammation [10]. Additionally, it has been demonstrated that the selective elimination of SCs increases both lifespan and healthspan in murine models [11].

In this study, we identified FDA-approved and experimental drugs as novel senolytics with a multi-target mechanism of action, capable of effectively inhibiting multiple SCAPs simultaneously. The molecules were identified through computational methods, including cheminformatic analysis, structure-based virtual screening, and molecular dynamics. We validated the senolytic activity of these molecules in two senescence models using primary cultures of human lung epithelial cells from the CCD8-Lu cell line. Specifically, two different models of cellular senescence induction were used: senescence induced by stress with H_2_O_2_ (stress-induced premature senescence, SIPS) and replicative senescence (RS).

## Results

### Structural Similarity Search

To identify new molecules with potential senolytic activity, we used structural similarity searching with 2D molecular fingerprints (FP) [12]. This simple yet powerful approach is based on the structure-activity relationship (SAR) concept, which allows for selecting and refining the most suitable drug candidates. This method assumes that molecules with similar chemical structures exhibit similar physicochemical and biological properties [13].

After evaluating various FPs (experimental section), we selected three of them to generate a consensus similarity to identify, within databases (DB) of FDA-approved drugs (FDA) and experimental drugs (EXP), the 100 drugs from each DB most structurally similar to any of the senolytics reported in the literature. Figure 1A shows the consensus structural similarity network, constructed with the MACCS keys 322, Atom Types, and EState Indices FPs, using the Tanimoto coefficient (T) as the similarity measure and a cutoff value of T > 0.7.

**Figure 1.**
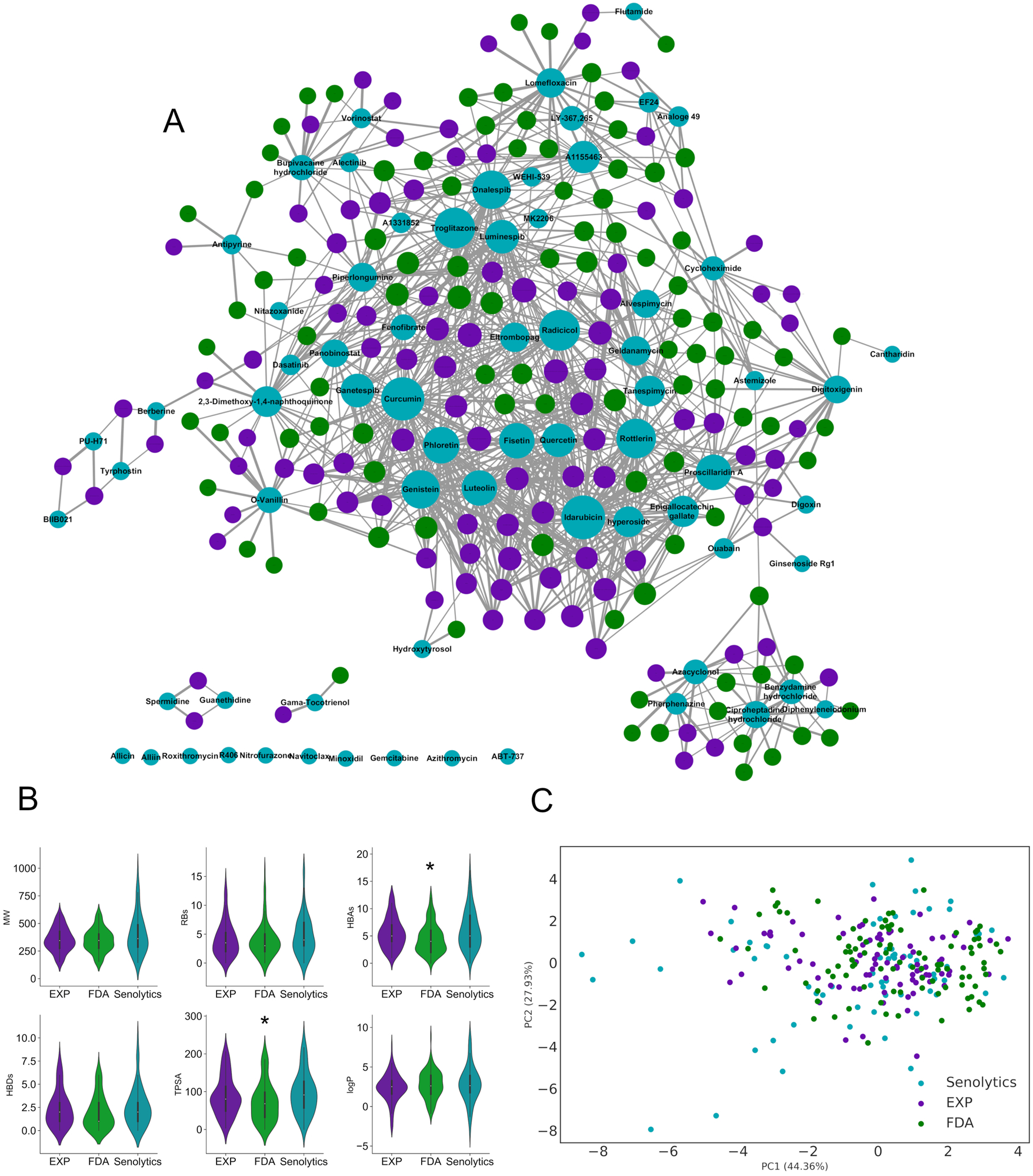
Structure Similarity Search. **A.** Consensus structural similarity network of the 200 drugs (100 FDA-approved and 100 experimental) most similar to senolytics constructed with the MACCS keys 322, Atom Types, and EState Indices fingerprints using a Tanimoto cutoff > 0.7. **B.** Statistical analysis of six molecular descriptors commonly used in medicinal chemistry for the selected 100 FDA-approved and 100 experimental drugs and senolytics. *p < 0.05*, Kruskal-Wallis test, post-hoc Dunnett with Holm adjustment. **C.** Exploration of the chemical space covered by senolytics and the selected FDA-approved and experimental drugs. The first two principal components retain 72.3% of the variance

In the network of the 100 molecules most similar to any senolytic from each DB, the turquoise blue nodes represent known senolytics, and their size is the number of molecules from the FDA and EXP DBs with which they share similarity. Green nodes represent molecules from the FDA DB, while purple nodes represent those from the EXP DB. Even at this relatively high cutoff point and considering the consensus, there are several drugs with a chemical structure similar to more than one senolytic.

Using the 100 molecules most similar to senolytics from each DB, we calculated and statistically analyzed six molecular descriptors commonly used in medicinal chemistry, as shown in Figure 1B, to verify if the SAR principle holds: molecular weight (MW), number of rotatable bonds (RBs), hydrogen bond acceptors (HBAs), hydrogen bond donors (HBDs), topological polar surface area (TPSA) and the octanol/water partition coefficient (logP). Only the FDA molecules show slight differences in the descriptors, having a lower number of HBAs (*p = 0.013*) and a slightly lower TPSA (*p = 0.029*) than the known senolytics.

Despite these slight differences, the 200 molecules filtered by similarity spanned a chemical space very similar to that of senolytics, as shown in Figure 1C, where the first two principal components of the principal component analysis (PCA) retain 72.3% of the variance. Taken together, these results suggest that the structural similarity search has detected 200 molecules with chemical structures similar to more than one known senolytic. Additionally, these molecules present physicochemical properties very similar to senolytics and hold a very similar chemical space, partially fulfilling the SAR principle, which could mean that they would also have a senolytic effect.

### Identification of Drug Targets

Once the molecules structurally similar to senolytics were identified, we determined the drug targets for these molecules. Gene expression data from human lungs generated by the GTEx project were used based on the premise that SCs accumulate in the body with increasing age, which allows for detecting changes in the expression levels of SCAPs in lung samples from elderly patients. The differential expression of 19 SCAPs, whose inhibition with shRNA produces selective death of SCs, was analyzed [4]. As shown in Figure 2, the expression levels of SCAPs in the elderly (60-70 years old) were compared to those in young adults (20-39 years old), quantifying the log2 of the fold change (FC). 11 SCAPs were found to have significant changes in their expression levels, of which only eight increased during aging. Four of these were selected: SERPINE1 (FC = 1.12, *p = 0.0001*), EFNB1 (FC = 1.04, *p = 0.005*), PDGFB (FC = 1.12, *p = 0.006*) and PI3KCD (FC = 1.12, *p = 0.01*). These four SCAPs were chosen because they are overexpressed and are part of a co-expression network that is enriched in protein-protein interactions in the human lung [14], suggesting that their inhibition could affect the formation of anti-apoptotic resistance networks.

**Figure 2.**
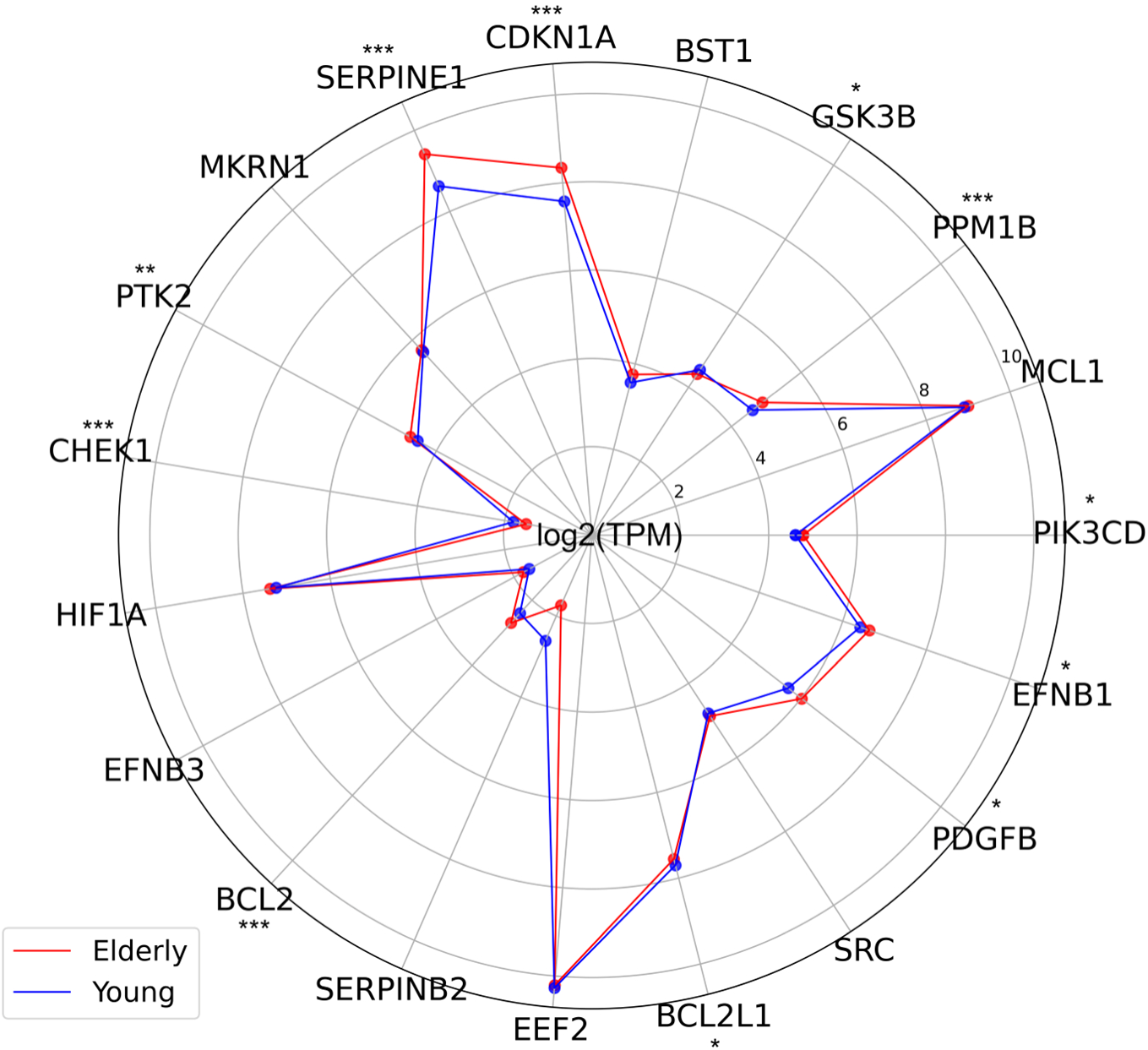
Differential Expression of 19 SCAPs. The expression of SCAPs was compared in human lung tissue from elderly (60-70 years, n=212) vs young (20-39 years, n=73) individuals. *\*p < 0.05*, *\*\*p < 0.005*, *\*\*\*p < 0.0005*, Wilcoxon test.

### Structure-Based Virtual Screening

After identifying the 200 molecules chemically most similar to senolytics, a consensus virtual screening was conducted using AutoDock Vina and AutoDock4 to target the three-dimensional structures of the proteins encoded by the four selected SCAPs. The goal was to identify molecules capable of binding simultaneously to the active sites of more than one target. The initial approach involved blind molecular docking. Figure 3 shows the results of the consensus virtual screening (CVS) for the EFNB1, PDGFB, SERPINE1, and PIK3CD proteins, where the x-axis represents the ranking of each molecule obtained in Vina. while the y-axis represents the ranking for AutoDock. The color gradient represents the average binding energy (kcal/mol) or affinity from both programs.

**Figure 3.**
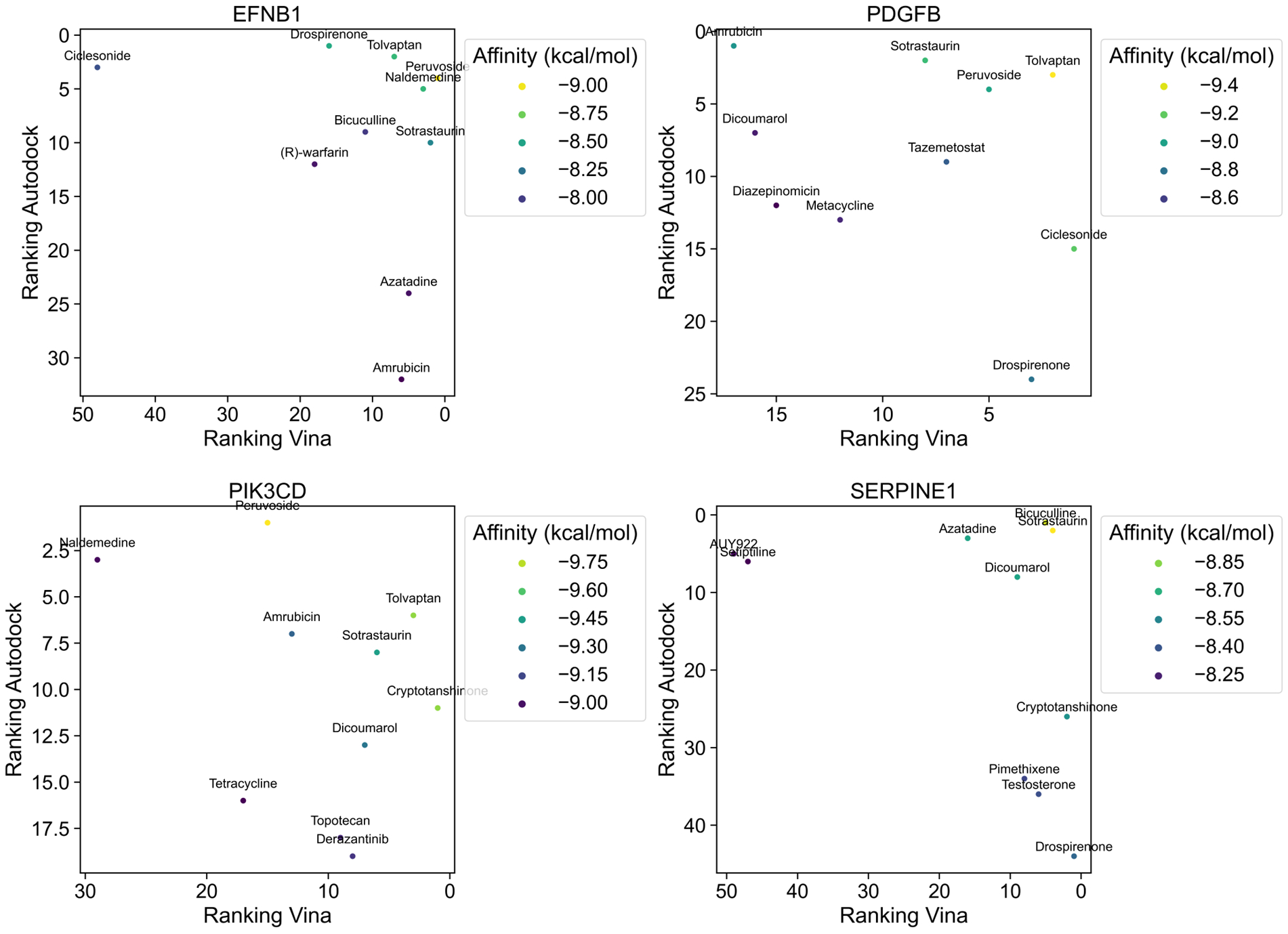
Consensus Virtual Screening. Molecular docking calculations of 200 molecules were performed on the entire 3D structure of the four SCAPs. The x-axis shows the rank obtained in AutoDock Vina, while the y-axis shows the rank of each molecule obtained in Autodock4. The color gradient shows the mean binding energy of AutoDock Vina and Autodock4. The ten molecules with the best consensus rank for each protein are shown.

Consequently, the molecules in the top right quadrant have the best ranking and binding energy in both molecular docking programs. Interestingly, we identified several molecules within the top 10 that present a good binding energy with more than one protein. Due to commercial availability and cost, we selected only four molecules whose chemical structure, potential targets, and current uses are summarized in Table 1; the rest of the multi-target molecules are shown in Supplementary Table S4, Supplementary File S1.

**Table 1.**
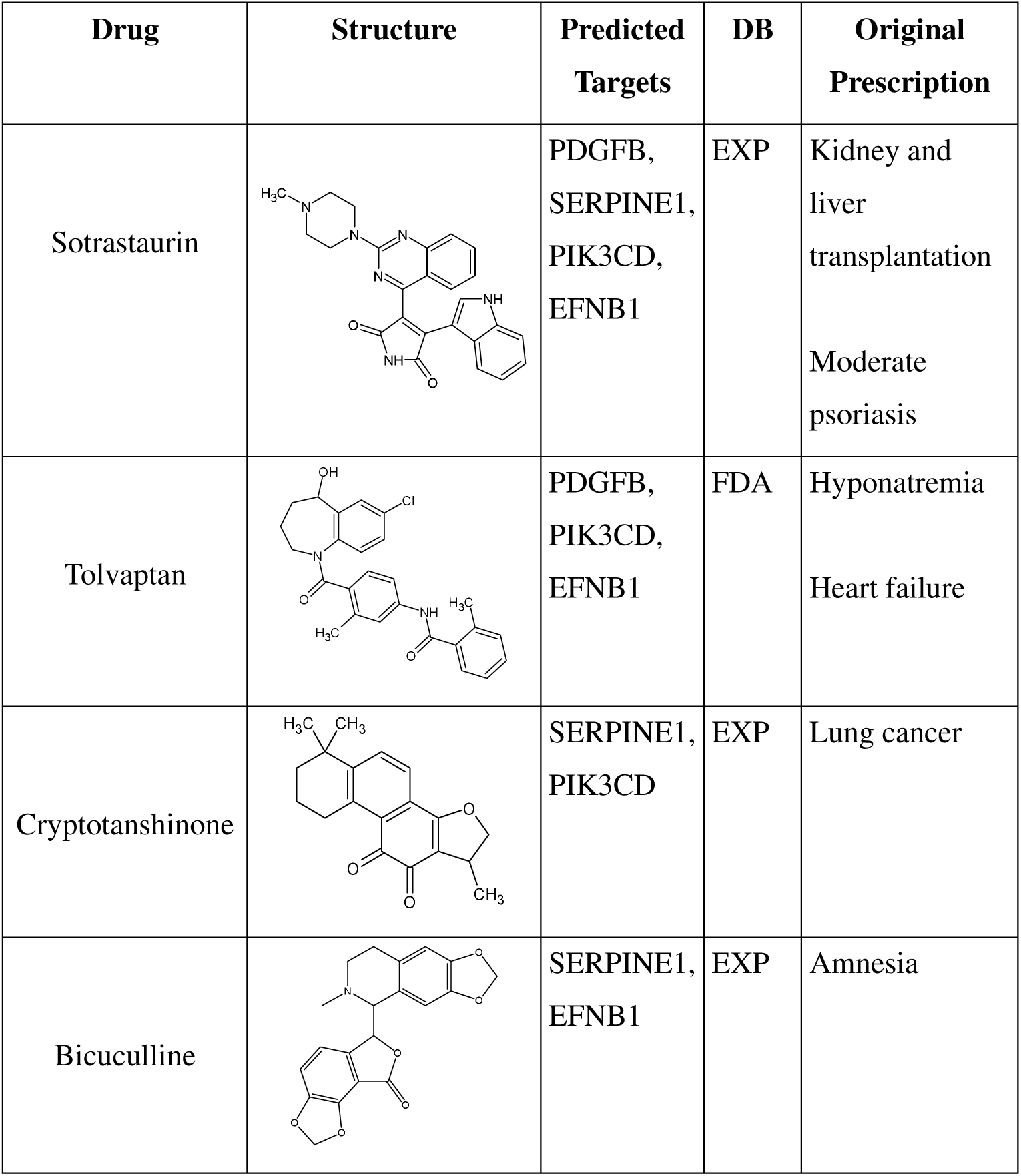
Drugs selected as computational hits. Four drugs were selected based on their multi-target potential, commercial availability, and cost.

| Drug | Structure | Predicted Targets | DB | Original Prescription |
| --- | --- | --- | --- | --- |
| Sotrastaurin |  | PDGFB,<br>SERPINE1,<br>PIK3CD,<br>EFNB1 | EXP | Kidney and liver transplantation<br>Moderate psoriasis |
| Tolvaptan |  | PDGFB,<br>PIK3CD,<br>EFNB1 | FDA | Hyponatremia<br>Heart failure |
| Cryptotanshinone |  | SERPINE1,<br>PIK3CD | EXP | Lung cancer |
| Bicuculline |  | SERPINE1,<br>EFNB1 | EXP | Amnesia |

### Docking on the Binding Site

Once the molecules with potential multi-target mechanisms were identified, the molecular docking calculations were refined by directing the molecules toward the binding site of each protein. First, we validated the molecular docking protocol by re-docking the co-crystallized inhibitors with the 3D structures, as shown in Supplementary Figure S4.

The results of the consensus docking using Vina and Autodock directed toward the binding site of SERPINE1, PIK3CD, PDGFB, and EFNB1 are shown in Figure 4. For each calculation, the results of the candidate molecules were compared with those of the co-crystallized inhibitor. This comparison was performed for all proteins except EFNB1, which lacks known inhibitors. Ideally, the candidate molecules should maintain most of the interactions the co-crystallized inhibitor shows and exhibit more negative binding energies than the co-crystallized inhibitor. These results were obtained for most of the ligand-receptor complexes, suggesting that these molecules could bind with good affinity to the binding site of more than one protein.

**Figure 4.**
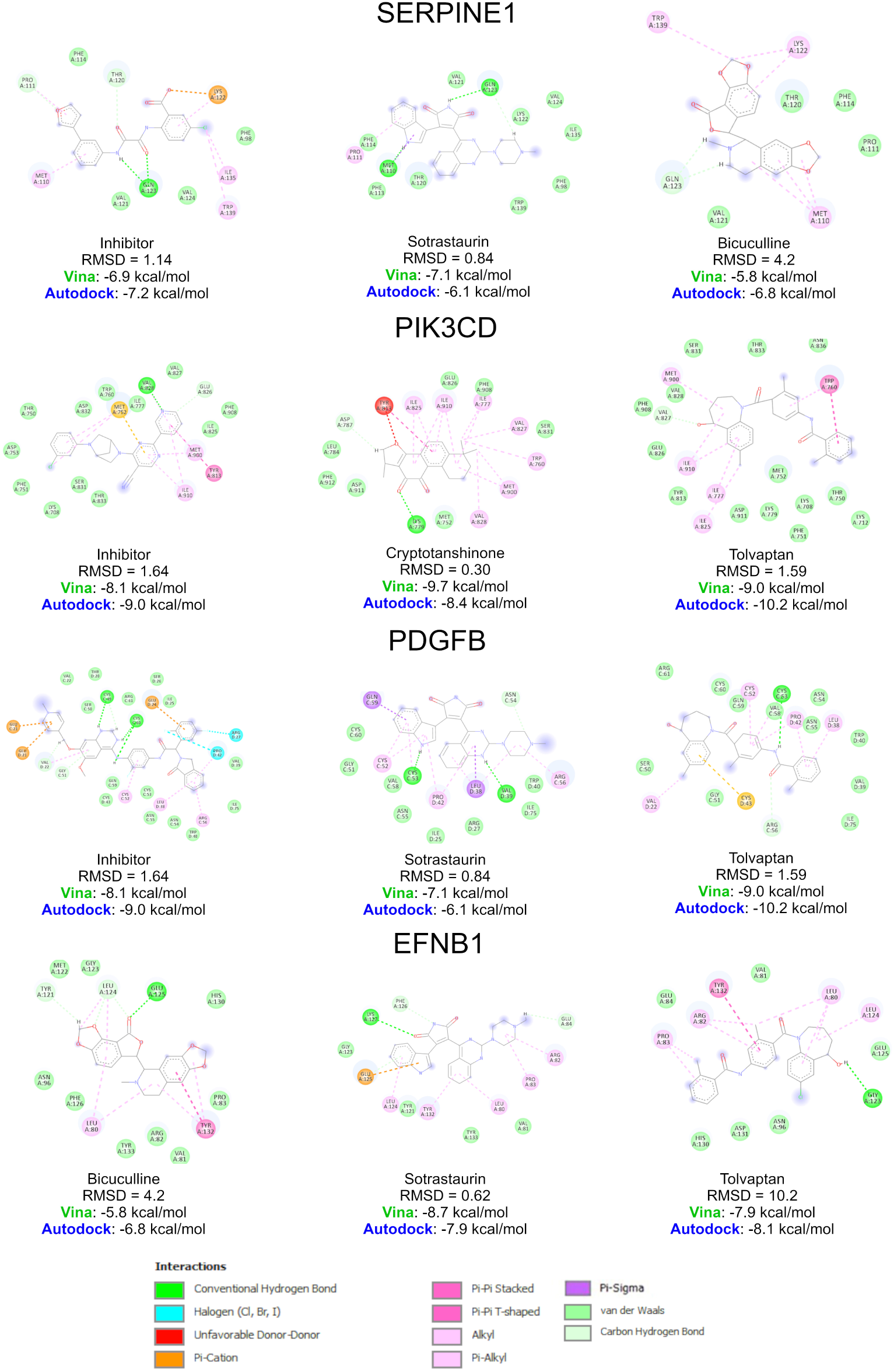
Molecular docking on the binding site. Using AutoDock Vina and Autodock4, docking calculations were performed on the binding site of each of the four proteins. The binding energy determined by each program and the RMSD between each program’s most energetically favorable positions are shown.

### Molecular Dynamics Simulations

The ligand-receptor complexes generated by AutoDock Vina underwent 100 nanoseconds (ns) of molecular dynamics simulations using the *YASARA Structure* software. Each drug was simulated with its respective potential target. Additionally, simulations were performed for the proteins in their apo state (without ligand) and with the co-crystallized or literature-reported inhibitors. Properties such as the root mean square deviation (RMSD) of the α-carbons of the proteins and the binding energy of the ligands were analyzed. A steered molecular dynamic (SMD) simulation was also carried out to evaluate the residence time of the ligand in the binding site under the application of an external force field.

As shown in Figure 5, the RMSD of the four proteins (EFNB1, PDGFB, PIK3CD, and SERPINE1) was compared in their apo form and in complex with various ligands. For PDGFB, PIK3CD, and SERPINE1, simulations were performed with their respective inhibitors (co-crystallized or reported in the literature). Additionally, the following candidate drugs were simulated: sotrastaurin (with EFNB1, PDGFB, and SERPINE1), bicuculline (with EFNB1 and SERPINE1), tolvaptan (with EFNB1, PDGFB, and PIK3CD), and cryptotanshinone (with PIK3CD). The RMSD analysis was conducted over 100 ns of simulation time for each system, providing insights into the structural stability of the proteins under different conditions.

**Figure 5.**
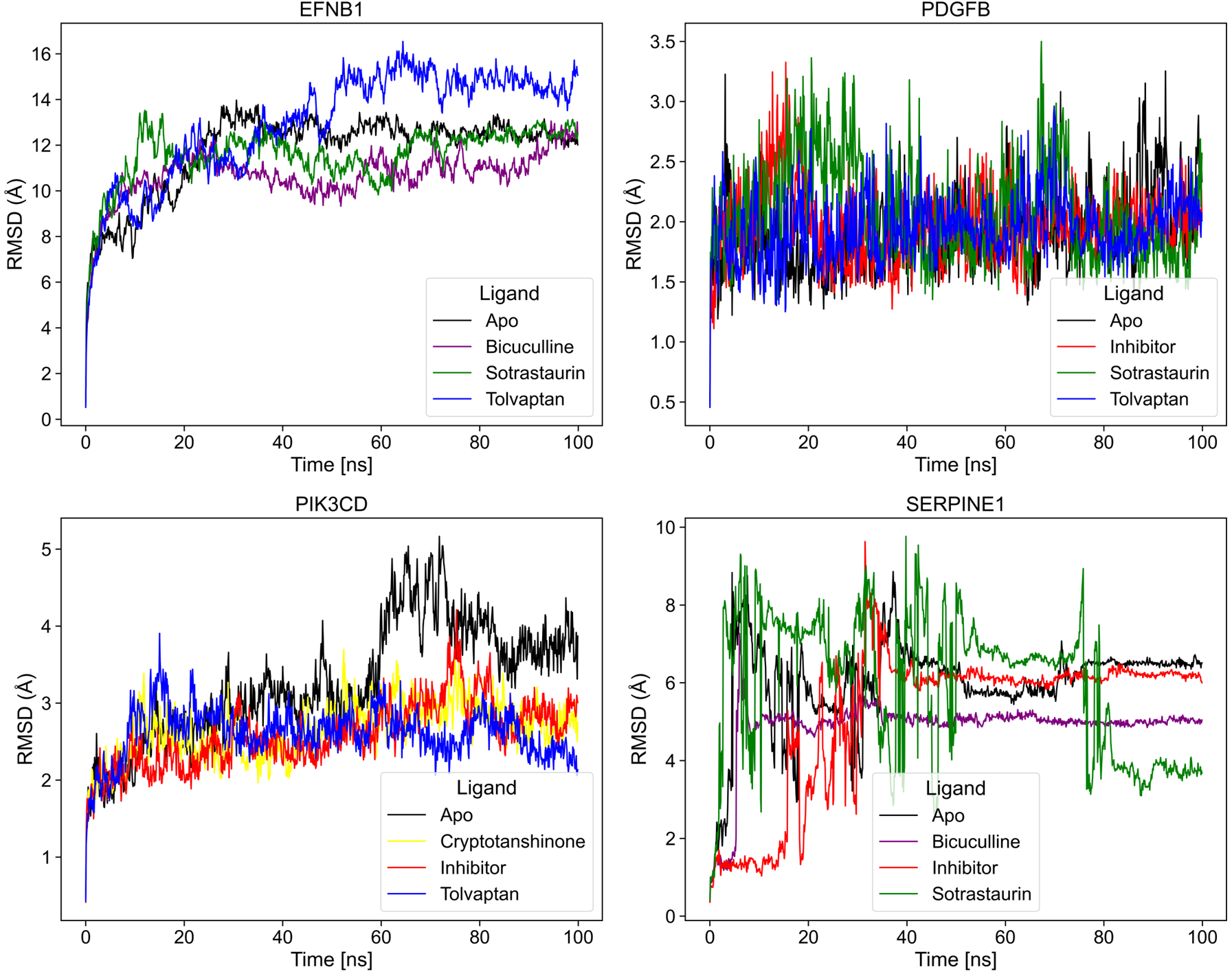
Molecular Dynamics Simulations. Analysis of the RMSD of the α carbons. The RMSD analysis of the α carbons was performed during 100 ns of simulation for the four proteins. For each protein, its Apo form (without ligands) was simulated. For PDGFB, PIK3CD, and SERPINE1, a co-crystallized or literature-reported inhibitor was simulated, and the corresponding candidate drugs were simulated.

For EFNB1, it was observed that the apo form stabilizes around 25 ns, with an average RMSD (aRMSD) = 11.79 Å and fluctuations below 2 Å, suggesting that this could represent its biologically active conformation. Sotrastaurin (aRMSD = 11.59 Å) and bicuculline (aRMSD = 10.59 Å) stabilize EFNB1 around 20 ns, maintaining an RMSD slightly lower than that of the apo conformation, indicating a possible stabilization of the protein structure. In contrast, tolvaptan (aRMSD = 12.95 Å) reaches stabilization at approximately 45 ns but with an RMSD higher than that of the apo form, which could indicate a conformational change that may affect the function of EFNB1.

The results for PDGFB indicate that the apo form (aRMSD = 1.91 Å) remains relatively stable throughout the simulation until the last 20 ns, where a slight increase in RMSD is observed. The inhibitor (aRMSD = 1.97 Å) shows similar stability to the apo form and stabilizes PDGFB from approximately 40 ns onwards for the rest of the simulation. Tolvaptan (aRMSD = 1.95 Å) and sotrastaurin (aRMSD = 2.09 Å) showed similar behavior to the inhibitor, stabilizing the protein around 35 ns and maintaining stability without abrupt peaks.

The RMSD values for PI3KCD suggest that the apo form (aRMSD = 3.26 Å) is the most unstable, showing increasing RMSD values over time. Both the inhibitor (aRMSD = 2.59 Å) and cryptotanshinone (aRMSD = 2.64 Å) exhibit similar behavior, stabilizing the protein with low RMSD values compared to the apo form. Interestingly, tolvaptan (aRMSD = 2.61 Å) maintains a behavior similar to the inhibitor throughout the simulation, with all three ligands showing more stable RMSD values than the apo form.

Finally, the results for SERPINE1 show the apo form (aRMSD = 6.0 Å) is highly unstable, suggesting the need for a conformational change for its biological activity. After 50 ns, the protein remains relatively stable. The inhibitor (aRMSD = 5.18 Å) starts the simulation with lower RMSD values. It maintains them until approximately 35 ns, at which point it adopts a similar behavior to that of the apo form but maintains lower RMSD values. Sotrastaurin (aRMSD = 5.96 Å) shows high instability throughout the simulation, with RMSD changes of up to 6Å in less than ten ns, suggesting that it fails to stabilize SERPINE1. On the other hand, bicuculline (aRMSD = 4.87 Å) stabilizes SERPINE1 around ten ns and keeps the protein stable with an RMSD lower than the inhibitor for the rest of the simulation.

### Binding Energy Analysis

The analysis of binding energies was performed using the MM-PBSA (Molecular Mechanics Poisson-Boltzmann Surface Area) method with *YASARA software*. This approach allows for a dynamic assessment of binding affinity over time, as shown in Figure 6. The binding energy (kcal/mol) was analyzed for co-crystallized inhibitors and four proposed senolytic drugs across four protein targets: EFNB1, PDGFB, PIK3CD, and SERPINE1. First, the energy of the complete ligand-protein complex system was evaluated in its bound state. Then, the energy of the system when the ligand and protein are separated at an infinite distance, representing the unbound state, was subtracted from this value. This process was repeated every 100 picoseconds throughout the 100 ns simulation. Within the AMBER14 force field framework, more negative values from this calculation indicate a stronger binding between the ligand and the protein [15].

**Figure 6.**
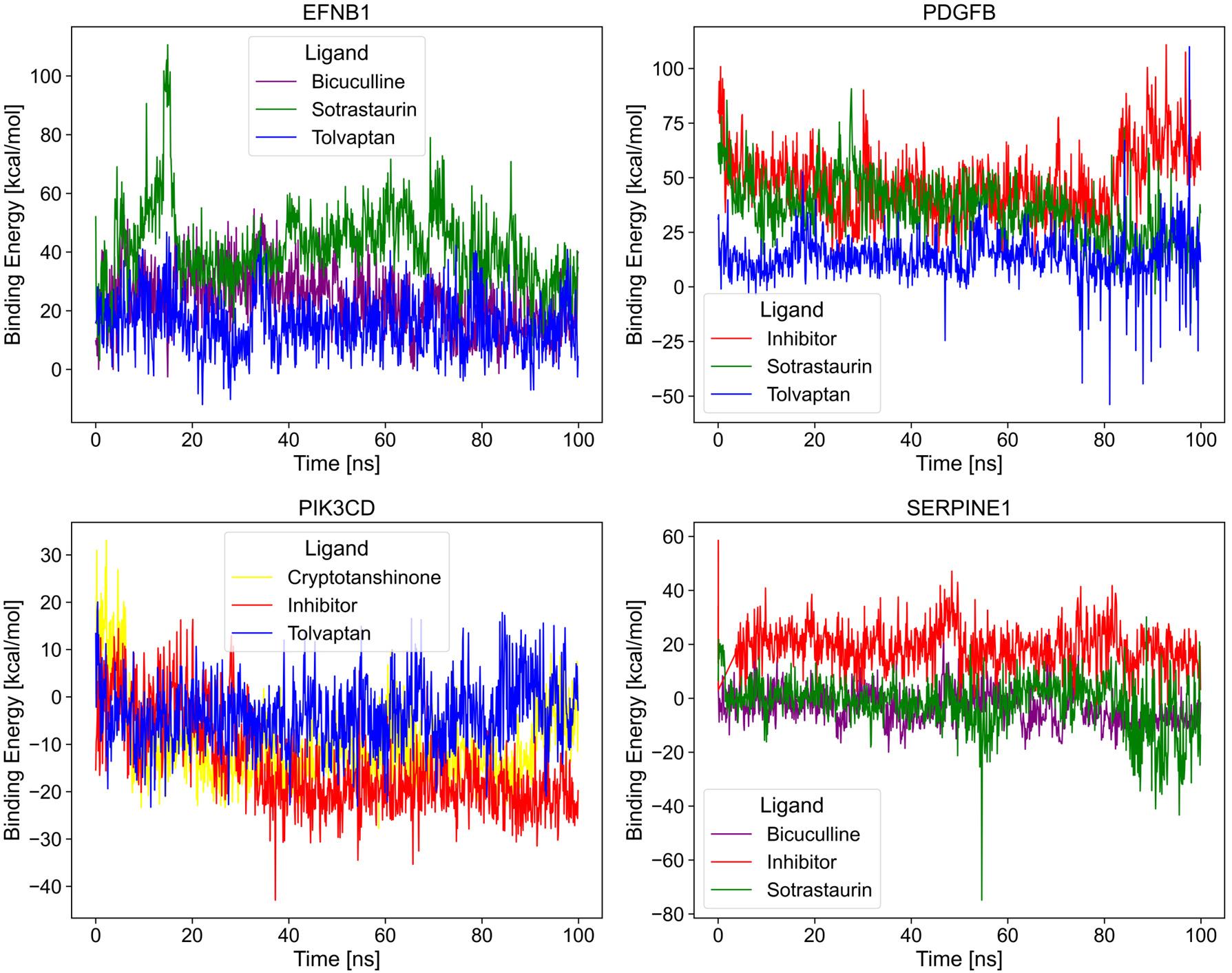
Binding energy analysis. Analysis of ligand affinity using the MM-PBSA method. The binding energy (kcal/mol) was analyzed over the 100 ns simulation time for the co-crystallized inhibitors and the four proposed senolytic drugs. Lower (more negative) binding energies indicate stronger affinity for the receptor.

For EFNB1, tolvaptan and bicuculline show relatively stable energies throughout the simulation. Tolvaptan generally exhibits lower binding energies, suggesting a better affinity for the protein than bicuculline. With average energies of 15.3 kcal/mol for tolvaptan and 23.55 kcal/mol for bicuculline. On the other hand, sotrastaurin exhibits larger energy fluctuations, with higher binding energies, indicating lower affinity with an average energy of 41.55 kcal/mol.

The results for PDGFB show that the inhibitor has the highest and most fluctuating positive energy, with an average of 47.67 kcal/mol. Tolvaptan and sotrastaurin maintain relatively stable energies for most of the simulation until 80 ns but exhibit abrupt peaks in the last 20 ns, with tolvaptan occasionally reaching very low binding energies, particularly in the latter part. The average energies for tolvaptan and sotrastaurin are 13.87 kcal/mol and 35.4 kcal/mol, respectively. This suggests that tolvaptan has a higher affinity for the receptor than the inhibitor.

For PIK3CD, all three molecules (cryptotanshinone, inhibitor, and tolvaptan) maintain stable and mostly negative energies throughout the simulation. The inhibitor consistently shows the lowest binding energies, followed by cryptotanshinone and tolvaptan. The inhibitor has an average energy of -14.29 kcal/mol, cryptotanshinone -10.38 kcal/mol, and tolvaptan -3.71 kcal/mol. This suggests that both tolvaptan and cryptotanshinone exhibit affinity for PIK3CD, with cryptotanshinone showing behavior more similar to the inhibitor.

Finally, the binding energies for SERPINE1, the inhibitor, exhibit generally higher binding energies compared to sotrastaurin and bicuculline, with an average of 19.4 kcal/mol. Sotrastaurin and bicuculline have average energies of -19.6 kcal/mol and -3.58 kcal/mol, respectively, suggesting that both drugs interact better with SERPINE1 than the co-crystallized inhibitor. Sotrastaurin shows periods of very low binding energies, particularly towards the end of the simulation, suggesting it may interact better with SERPINE1 compared to the co-crystallized inhibitor and bicuculline.

### Steered Molecular Dynamics (SMD)

SMD is a variant of traditional molecular dynamics. Its goal is to mimic experimental conditions that cannot be easily reproduced in standard molecular dynamics simulations. The main feature of SMD is the ability to apply external forces to “steer” the system along a specific trajectory or towards a particular conformational state. For example, it can be used to extract a ligand from its binding site in a protein, providing valuable information about the forces and energies involved in these processes, as shown in Figure 7.

**Figure 7.**
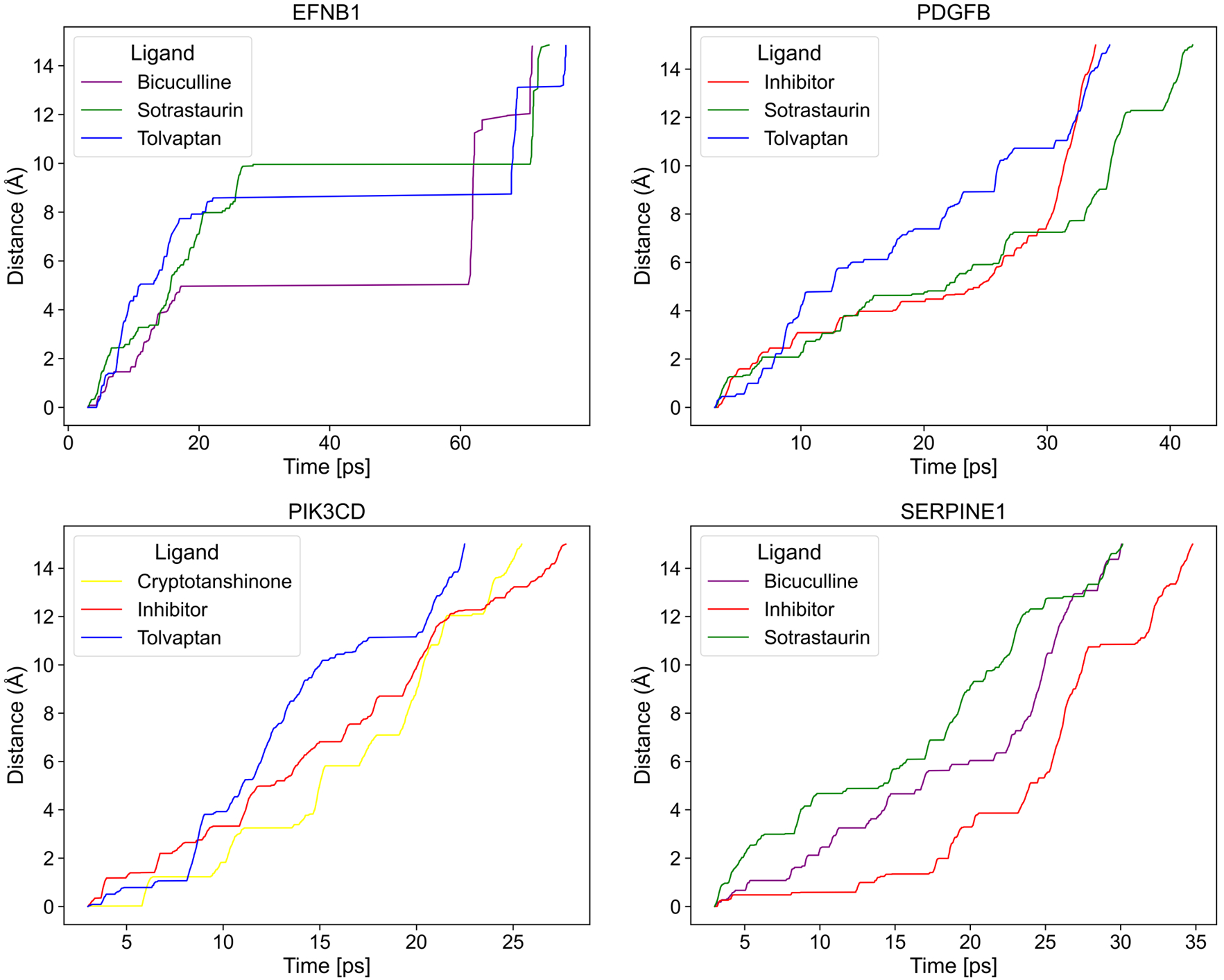
Steered Molecular Dynamics Simulations. The time it takes for each drug to exit the binding site when subjected to an external force field and constant velocity was quantified. The ligand distance over time is compared against the known inhibitor.

The SMD results for EFNB1 are shown in the top left panel of Figure 7. It is observed that tolvaptan, bicuculline, and sotrastaurin have a similar residence time in the binding site of around 70 ns.

Additionally, all three drugs have very pronounced plateaus, meaning they remained in the same site for an extended time, even as the external force field increased, suggesting a good affinity for the receptor. Notably, bicuculline demonstrates the most pronounced plateau, indicating potentially stronger binding. In the top right panel, the results for PDGFB are shown. The inhibitor leaves the binding site around 32 ns, although tolvaptan exhibits higher distances from the binding site than the inhibitor when the same force is applied. The time it took to leave the active site is slightly longer than that of the inhibitor. Sotrastaurin took around 43 ns to leave the binding site; these results suggest that both tolvaptan and sotrastaurin have good affinity for PDGFB.

In the bottom left panel, the results for PIK3CD are shown. The inhibitor leaves the active site around 28 ns, while cryptotanshinone exhibits behavior similar to the inhibitor’s, leaving the active site at 25 ns. Tolvaptan maintains the highest distances and has the shortest abandonment time of 22 ns. Finally, the results for SERPINE1 are shown in the bottom right panel. The abandonment time of the inhibitor was 35 ns, while sotrastaurin and bicuculline, both with very similar behavior, leave the binding site at 30 ns.

### *In Vitro* Senolytic Effect

To evaluate the senolytic effect of the computationally identified molecules, the MTT assay was used, and their impact was examined in two senescence models that we had previously reported [14], the stress-induced premature senescence (SIPS) and the replicative senescence (RS). In this study, primary human lung fibroblasts from the CCD8-Lu cell line were used.

Figure 8 shows the results of the MTT assay for 48 hours of senolytic treatment. The control groups’ relative optical density (ROD) (senescent cells and proliferating cells without treatment) was normalized to 1, and the other groups were normalized to their respective untreated control. Statistical analysis was performed using a one-way ANOVA and a Tukey post-hoc test with Benjamini-Hochberg adjustment to control the false positive rate. Panel A shows the results of the assays for the SIPS model. Cryptotanshinone exhibits a senolytic effect at 15 μM (*p = 0.0003*) and 20 μM (*p < 1×10^-17^*), and sotrastaurin has an effect at 30 μM (*p < 1×10^-17^*). At the same time, tolvaptan and bicuculline did not show a senolytic effect, even at concentrations of 40 μM. Interestingly, both cryptotanshinone 20 μM (*p < 1×10^-17^*) and sotrastaurin 30 μM (*p = 0.003*) had a greater senolytic effect than the reference combination D+Q (250 nM and 50 μM, respectively).

**Figure 8.**
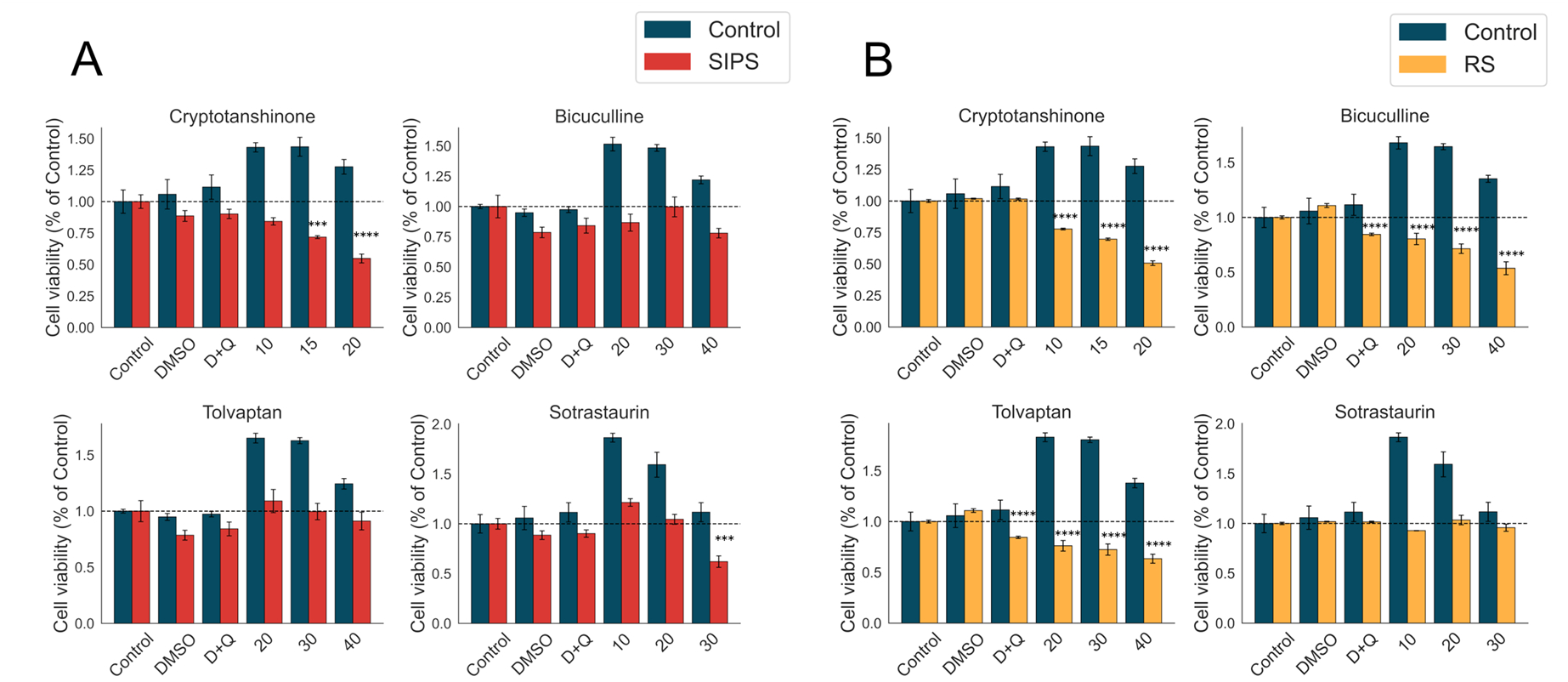
In vitro evaluation of the senolytic effect at 48 hours. **A.** MTT assay in the SIPS model. Cryptotanshinone shows a senolytic effect at 15 μM and 20 μM, while sotrastaurin has an effect at 30 μM. Tolvaptan and bicuculline do not exhibit a senolytic effect. **B.** MTT assay in the RS model. Cryptotanshinone has a concentration-response effect starting at 10 μM. Tolvaptan and bicuculline exhibit a senolytic effect in RS, with a concentration-response effect starting at 20 μM. *\*\*p ≤ 0.001*, *\*\*\*p < 0.0001*, *\*\*\*\* p < 0.00001*.

**Figure 9.**
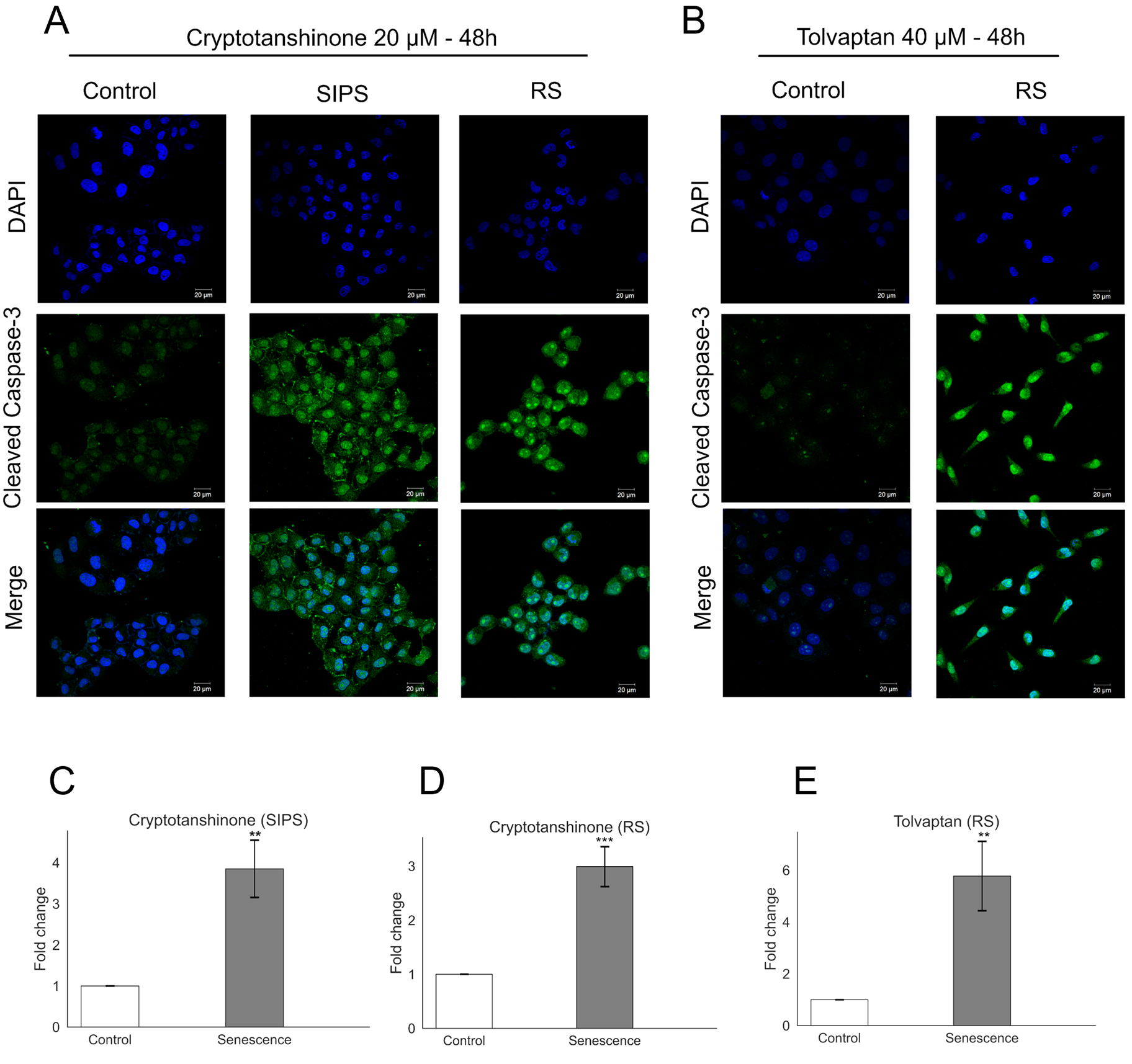
Selective apoptosis induction. **A.** IF showing the expression of active Caspase-3 in cells treated with 20 μM cryptotanshinone for 48 hours in the SIPS model. **B.** IF showing the expression of active Caspase-3 in cells treated with 40 μM tolvaptan for 48 hours in the RS model. **C.** Quantification of active Caspase-3 expression in the SIPS model, showing a significant ∼4-fold increase in SCs treated with cryptotanshinone. **D.** Quantification of active Caspase-3 expression in the RS model, showing a significant ∼3-fold increase in SCs treated with cryptotanshinone. **E.** Quantification of active Caspase-3 expression in the RS model, showing a significant ∼6-fold increase in SCs treated with tolvaptan. *\*\*p ≤ 0.001*, *\*\*\*p < 0.0001*.

To corroborate the senolytic effect, the same experiments were performed in the RS model, as shown in panel B. Cryptotanshinone had a dose-response effect starting at 10 μM (*p < 1×10^-17^*), which increased for the 15 and 20 μM concentrations (*p < 1×10^-17^*). As observed in the SIPS model, sotrastaurin did not show a senolytic effect at 30 μM. Interestingly, tolvaptan and bicuculline did exhibit a senolytic effect on the RS model. Tolvaptan had a clear concentration-response effect that began at 20 μM (*p < 1×10^-17^*) and increased at the 30 and 40 μM concentrations (*p < 1×10^-17^*). Bicuculline showed the same concentration-response behavior starting at 20 μM (*p < 1×10^-17^*) and increased at the 30 and 40 μM concentrations (*p < 1×10-17*). In particular, in the last two experiments with cells at passage > 70 with D+Q, a senolytic effect was observed (*p < 1×10^-17^*); however, two of the candidate drugs had a greater senolytic effect at their lowest concentrations: cryptotanshinone 10 μM (*p < 1×10^-17^*) and tolvaptan 20 μM (*p = 0.035*), while bicuculline at 30 μM had a greater senolytic effect than the D+Q combination (*p = 0.0001*).

The MTT assay measures cell viability as a function of the redox activity of the cells, and this activity correlates with the amount of dead or live cells. However, the MTT assay does not indicate whether the apparent senolytic effect is due to the selective death of senescent cells. To verify that the tested drugs induced selective apoptosis, the expression of the active Caspase-3 protein was evaluated by immunofluorescence in the SIPS model. As shown in panel A, increased expression of active Caspase-3 is observed in SCs treated with 20 μM cryptotanshinone for 48 hours; the expression of this protein is an indicator of the execution of apoptosis. The quantification of active Caspase-3 expression shows an almost 4-fold increase (*p = 0.001*) of this protein in senescent cells compared to proliferating cells (Panel C), confirming that the decrease in cell viability detected by the MTT assay is due to the selective induction of apoptosis.

Additionally, it was analyzed whether the decrease in viability observed in the RS model was also due to the induction of selective apoptosis in senescent cells. As shown in Panel B, both cryptotanshinone 20 μM and tolvaptan 40 μM at 48 hours induced the selective death of senescent cells without affecting proliferating cells. As shown in Panels D and E, the quantification of pixel density indicates that the activation of active Caspase-3 is almost three times higher in senescent cells when treated with cryptotanshinone (*p = 0.0003*) and almost six times higher with tolvaptan (*p = 0.001*).

## Discussion

The elimination of SCs through senolytics could stop or reverse various chronic-degenerative diseases associated with aging. However, to date, there is contradictory evidence in this regard, as eliminating certain types of SCs could accelerate or promote the onset of pathologies since some types of SCs may be necessary to maintain the healthspan [16]. It has been shown that eliminating SCs using senolytics could negatively affect regeneration [17] and tissue architecture [18] due to the reduced appearance of regenerative epithelial stem cells. The aforementioned issues could be largely resolved by adopting rational drug design approaches [19] in senotherapy since most of the senolytics discovered to date have been identified through classical and agnostic target methods, such as massive phenotypic screening [20], [21], hypothesis-based methods [4], [22], or by serendipity [23].

To increase the probability of identifying novel senolytics, we used a rational design approach focused on polypharmacology. This concept refers to the design or use of drugs that act on multiple targets or biological pathways simultaneously. Instead of the traditional “one drug, one target” strategy, polypharmacology recognizes that many diseases and complex phenotypes are multifactorial and can be addressed more efficiently with drugs that have multiple therapeutic effects through interaction with various molecular targets [24].

We used gene expression data from human lungs to identify the drug targets to direct molecules with potential senolytic activity. CS and the discovery of senolytics have been widely studied in various human lung fibroblast cell lines such as WI-38 [25] and IMR-90 [22]. Both cell lines originate from newborns younger than ten months, whereas CCD-8Lu is derived from an adult. Adult cells could provide information closer to the accumulation of damage and response to drugs compared to newborns. It was assumed that during aging, more SCs accumulate [26] and that using transcriptomic data, we could identify differentially expressed SCAPs in this tissue, thus directing the molecules against biologically significant survival targets. However, this approach has some weaknesses. It is known that the levels of messenger RNA expression are not correlated with the expression levels of the proteins they encode [27]. Furthermore, the development and activation of SCAPs are highly dynamic and could depend on the cell type, the senescence-inducing stimulus, and the senescence stage, as occurs with SASP [28].

The four selected SCAPs were: 1) Platelet-Derived Growth Factor B (PDGFB). PDGFB binds to PDGF receptors on the cell surface, triggering a signaling cascade that leads to cell proliferation and survival. PDGFB promotes cell survival by activating the PI3K/AKT signaling pathway, inhibiting apoptosis through phosphorylation and inactivation of pro-apoptotic components such as BAD and the upregulation of anti-apoptotic proteins [29]. 2) Phosphatidylinositol 3-Kinase, Delta Subunit (PIK3CD), which is a catalytic subunit of the class IA phosphatidylinositol 3-kinase (PI3K). Its activation produces PIP3, a second messenger that activates AKT and other signaling pathways. Activated AKT can phosphorylate and neutralize various pro-apoptotic effectors, promoting cell survival [30]. 3) Plasminogen Activator Inhibitor-1 (SERPINE1), an inhibitor of proteases that regulates the activity of proteases such as plasminogen and urokinase. It can interact with integrins and other receptors on the cell surface, activating signaling pathways that promote survival. It has been associated with resistance to apoptosis in cancer cells as it can inhibit Caspase-3 [31]. 4) Ephrin-B1 (EFNB1), a member of the ephrin family of proteins that binds to Eph receptors. EFNB1 interacts with EphB receptors, triggering signaling cascades that affect cell adhesion, shape, and mobility, promoting survival and migration of some tumor cells [32].

Our method identified four drugs that could potentially inhibit more than one SCAPs. None of these drugs have been reported as senolytic to date. However, it’s interesting to note that the method identified experimentally validated inhibitors of SCAPs. One of the most interesting cases is sotrastaurin, which our screening identified as a potential inhibitor of PIK3CD. However, experimental evidence has shown that it does not have an affinity for that specific target. Nevertheless, it has been validated as an inhibitor of other SCAPs, including KDR, GS3KB, EPHA2, KIT, FLT1, ABL1, and SRC [33]. While sotrastaurin is not an inhibitor of PIK3CD, studies have shown that in combination with PI3K/AKT inhibitors, it enhances the inhibition of that signaling pathway [34]. On the other hand, tolvaptan has been demonstrated to inhibit the PI3K/AKT signaling pathway [35], [36], but no direct affinity assays have been performed on PIK3CD. Cryptotanshinone has been reported as an inhibitor of SERPINE1 through enzymatic assays [37] and has been shown to inhibit PI3K/AKT signaling [38].

Previously, we reported that the average IC_50_ value of senolytics is 15 μM [39]; therefore, we decided to work within a therapeutic window of 10 to 40 μM. A preliminary pilot assay (n = 1 independent experiment performed in triplicate; data not shown) of various concentrations served as a guide for choosing the best concentration range for each drug. The results of the MTT assays show that sotrastaurin at 30 μM reduced the functionality of cells in SIPS but not in RS. These results support the hypothesis of the differential development of SCAPs since, although sotrastaurin is an experimentally validated inhibitor of multiple SCAPs, it cannot reduce the viability of SCs. An interesting result was obtained with cryptotanshinone, which showed a reduction in viability in both senescence models at practically the same concentrations, slightly more potent in RS.

On the other hand, tolvaptan and bicuculline also showed an effect in the RS model but not in the SIPS model, reinforcing the hypothesis of the differential development of SCAPs. Furthermore, this result could suggest that some of the molecules have a higher affinity for certain receptors since, according to the computational results, cryptotanshinone, sotrastaurin, and bicuculline inhibit SERPINE1. The first two drugs had an effect in SIPS, but only cryptotanshinone had an effect in RS, while bicuculline did not have an effect in SIPS but did in RS, which would indicate that the expression of SERPINE1 increases in RS.

The D+Q combination, the first discovered and most widely studied senolytic, served as our positive control. This combination is currently in clinical trials for treating idiopathic pulmonary fibrosis [9] and diabetic renal complications [7]. Notably, our four newly identified molecules demonstrated superior senolytic activity compared to D+Q in both senescence models studied.

While D+Q inhibits the PI3K/AKT signaling pathway, its lack of senolytic activity in our models suggests this pathway may not be crucial for lung fibroblast survival in these specific senescence states. Consequently, our four molecules could offer safer and more effective therapeutic options, particularly for treating idiopathic pulmonary fibrosis. Considering these facts, any of the four molecules identified in this work could represent a safer and more effective therapeutic option, mainly for treating idiopathic pulmonary fibrosis. Tolvaptan, in particular, shows promise for treating diabetic kidney complications, given its established use in renal conditions [40].

Although this proposal may seem ambitious, our findings, as illustrated in Figure 7, support these newly discovered drugs’ improved efficacy and safety. The senolytic effect of our molecules was highly selective, inducing apoptosis exclusively in SCs. Remarkably, normal cells treated with these molecules showed increased viability, indicating low or no toxicity. This apparent proliferation boost in normal cells could be advantageous, potentially addressing the cell density reduction issue associated with SC removal.

## Conclusions

In this work, we report the senolytic activity of sotrastaurin, cryptotanshinone, tolvaptan, and bicuculline, which were identified through rational drug design approaches and the combination of various computational tools such as structural similarity search with molecular fingerprints, cheminformatics analysis, structure-based virtual screening, and molecular dynamics simulations. The senolytic activity of these four drugs is mediated by a multi-target mechanism through inhibiting multiple SCAPs such as SERPINE1, PIK3CD, PDGFB, and EFNB1. Additionally, cryptotanshinone demonstrated the ability to induce selective apoptosis of senescent cells under two different senescence-inducing stimuli.

Moving forward, exploring the senolytic effect in various cell types, including epithelial cells, endothelial cells, neurons, and astrocytes, under different senescence states, such as IRS, OIS, SIPS, and RS, represents a fundamental approach to understanding the versatility of senolytics and their application in a broad spectrum of conditions related to cellular senescence and aging. Furthermore, it is important to validate the potential ability of these senolytics to induce proliferation in normal cells. This could be achieved by quantifying the expression of proliferation markers, such as the Ki-67 nuclear proliferation antigen. If senolytics can stimulate proliferation in normal cells without inducing senescence, they could represent a valuable tool for tissue regeneration and overall health promotion.

Finally, given the context of the COVID-19 pandemic and the severe pulmonary sequelae experienced by some patients, conducting pilot clinical trials with FDA-approved drugs and those in the experimental phase would be highly beneficial. These trials should focus on patients presenting significant pulmonary complications due to COVID-19 infection. Evaluating the ability of these senolytics to improve lung function and reduce chronic inflammation could positively impact the quality of life of these patients. Our proposed future directions build upon our current findings and highlight the potential of our computational approach in identifying effective senolytic compounds for various applications in age-related diseases and for addressing emerging health challenges.

## Materials and Methods

### Molecular Filtering Rules

To build the different databases (except the reference database of senolytics), the following desirable physicochemical properties in pharmaceutical chemistry were used [41], [42]: molecular weight: 150–550 g/mol, ring systems: 0–10, number of carbons: 5–40, rotatable bonds: 0–15, allowed atoms: H, C, N, O, F, S, Cl, Br, and P, partition coefficient log P: between -0.4 and +5.6, should not contain more than ten hydrogen bond acceptors (nitrogen, oxygen or fluorine atoms).

### Molecular Databases

Senolytics: A database was manually generated by searching PubMed with the term “Senolytics”. 70 molecules were found until 20/02/2020. The database contains the molecule’s name, molecular weight, chemical structure, and reference. The SMILES (Supplementary Table S1, Supplementary Material S1) were also obtained from PubChem for each compound to generate an SDF file.

FDA-approved drugs (FDA): The Drugbank 5.1.6 database [43] was obtained, which originally contained 2,466 FDA-approved drugs. The SDF file was processed with *Datawarrior 5.0* [44], and after applying the established filtering rules, the database was reduced to 1,984 drugs.

Experimental drugs (EXP): The same Drugbank database and methodology were used for the FDA-approved drug database. The filtering reduced the database to 7,552 experimental drugs. The three databases can be consulted in the Supplementary Material S2 (Excel file).

### Structural Similarity

The structural similarity search was performed using FP calculated with the *Mayachemtools* package [45]. The senolytics database was used as a reference, and the FDA and EXP databases were used as search databases. Once the FP were generated, the similarity matrix was calculated using the Tanimoto coefficient [46]. Subsequently, the performance of the FP was evaluated using cumulative distribution function (CDF) curves of the similarity matrices (Supplementary Figure S1, Supplementary Material S1), and descriptive statistics were calculated (Supplementary Tables S2 and S3, Supplementary Material S1). Additionally, Pearson correlation matrices were constructed, and it was determined to use the MACCS keys 322, Atom types, and EState Indices FPs because they were poorly correlated and provided different structural information (Supplementary Figure S2, Supplementary Material S1). The filtering of the molecules with the highest structural similarity was performed by selecting the highest Tanimoto coefficient for each molecule for each of the three FPs. Subsequently, the mean of the maximum Tanimoto coefficients of the three selected FPs was calculated, the molecules were ranked, and the top 100 molecules from each database were selected. To visualize the similarity between the candidates and the senolytics, a consensus similarity network was constructed by averaging the Tanimoto coefficient value of the three FPs for each molecule and applying a cutoff of 0.7; this threshold has been previously reported by us as significant within the similarity of senolytics [39]. The visualization was performed with *Cytoscape 3.10.1.* The names of the 200 molecules most similar to senolytics and their consensus similarity values can be consulted in the Supplementary Material S2 (Excel file).

### Statistical Analysis of Molecular Descriptors

Using the *RDKIT* library [47], six molecular descriptors commonly used in medicinal chemistry [48] were calculated: Molecular Weight (MW), Number of Rotatable Bonds (RBs), Hydrogen Bond Acceptors (HBAs), Hydrogen Bond Donors (HBDs), Topological Polar Surface Area (TPSA), and the Octanol/Water Partition Coefficient (logP). The violin plots were constructed using the *Seaborn 0.10.1* library. A Shapiro-Wilk normality test was performed with the *Scipy 1.5.2* library using a significance level of *p < 0.05*. As normality was not met, Kruskal-Wallis was selected as the statistical test implemented with the *Scipy* library, followed by a Dunnett post-hoc test with Benjamini-Hochberg adjustment using the *Scikit 0.23.1* library.

### Chemical Space Exploration

The same 6 molecular descriptors previously calculated with *RDKIT* (MW, RBs, HBAs, HBDs, TPSA, and logP, Supplementary Material S2) were used. Dimensionality reduction was performed using PCA with the Scikit library. The molecules and their molecular descriptors for each database can be consulted in the Supplementary Material S2 (Excel file).

### Identification of Drug Targets

Gene expression data from human lung of men and women, normalized by transcripts per million (TPM), generated by RNA-Seq from the GTEx V8 project [49], were obtained. No other normalization or transformation was applied. The samples were divided into two groups: young (20– 30 years, n= 73) and elderly (60–79 years, n= 212). The expression values of 19 SCAPs were filtered and averaged, and fold changes (FC) were calculated using the formula: log2 (mean expression in elderly) / log2 (mean expression in young) for each gene. The statistical significance of the change in mean expression levels of each of the 19 SCAPs was determined using a Wilcoxon rank-sum test with the ranksums function from the Scipy library. This test is suitable for large datasets (n > 8), as it produces a low number of false positives [50].

### Virtual Screening

#### Protein Preparation

The three-dimensional structure from the Protein Data Bank (PDB) of the proteins PDGFB (ID: 4QCI), PIK3CD (ID: 6OCO), EFNB1 (ID: 6THG), and SERPINE1 (ID: 3UT3) was obtained. Protein processing was performed with the *UCSF Chimera software* 1.17.3 [51], which consisted of removing water molecules, removing bound ligands, assigning protons, and energy minimization. The chosen force field was AMBER FF14SB [52].

#### Ligand Preparation

The databases with the molecules were processed with *Open Babel Software 2.3.1*. [53]. The processing consisted of assigning protonation states and minimizing the structures. The chosen force field was MMFF94 [54].

#### Molecular Docking

For virtual screening, the 100 candidates with the highest similarity to any senolytic from the FDA and EXP databases were chosen. Due to errors in various structures the total number of molecules was 168. *Pyrx 0.8* [55] was used to perform the screening with *Autodock Vina 1.1.2* [56] with an exhaustiveness of 100 cycles. Additionally, *Autodock 4.2.6* [57] was used with medium exhaustiveness and a genetic algorithm. The docking for each molecule was blind, covering the entire processed protein.

#### Consensus

The consensus virtual screening was obtained by normalizing the data obtained from each molecular docking program using z-scores. Once the z-scores of each molecule and each pose were obtained, only the scores of the pose with the lowest binding energy were kept. These poses were summed for each program and ranked from highest to lowest z-score. The mean for both programs was obtained, and the list was ranked again from highest to lowest score.

### Molecular Docking on the Binding site of SCAPs

#### Target Processing

The processing of proteins and ligands was performed in the same way as for virtual screening. SERPINE1 active site:

The 7AQF structure from the PDB co-crystallized with an experimentally validated inhibitor (CHEMBL4210355) of SERPINE1 [58] was obtained, the active site is the following: Met^110^, Pro^111^, Thr^120^, Lys^122^, Gln^123^, Ile^135^, Trp^139^. Molecular docking parameters: Center X: 35.561, Y: -4.298, Z: - 0.660. Dimensions (Angstrom) X: 12.585 Y: 10.101 Z:11.617

##### PIK3CD active site

The 6OCO structure from the PDB co-crystallized with an experimentally validated inhibitor (M5V) of PIK3CD [59] was obtained, the active site is the following: Met^752^, Tyr^813^, Val^828^, Met^900^, Ile^910^. Molecular docking parameters: Center X: 39.024 Y: 13.576 Z:35.174 Dimensions (Angstrom) X: 14.423 Y: 9.405 Z:12.053

##### PDGFB active site

The 4QCI structure from the PDB was obtained, the reported active site [60] is the following: Leu^38^, Val^39^, Trp^40^, Gln^71^, Arg^73^, Ile^75^, Ile^77^, Arg^79^, Lys^80^, Lys^81^, Pro^82^, Ile^83^, Phe^84^, Lys^85^, Lys^86^. The experimentally validated inhibitor CHEMBL4443483 was obtained from ChEMBL. Molecular docking parameters: Center X: 20.116 Y: -1.392 Z:12.659. Dimensions (Angstrom) X: 19.423 Y: 11.848 Z: 22.419

##### EFNB1 active site

The 3D structure (AF-P98172-F1) from Alphafold [61] was obtained, the active site was determined using the online server Prankweb [62]. The identified active site is the following: Leu^124^, Gly^125^, Phe^126^, Tyr^132^, Leu^80^, Val^81^, Arg^82^, Pro^83^. Molecular docking parameters: Center X: -13.176 Y: 31.101 Z: -4.886. Dimensions (Angstrom) X: 19.816 Y: 21.747 Z: 16.584

For each protein, molecular docking was performed using *Autodock Vina 1.1.2* with an exhaustiveness of 100 cycles and *Autodock 4.2.6* with medium exhaustiveness using a genetic algorithm. The positions with the best affinity (Pose 0) for both programs were selected, and the RMSD between the two structures was calculated using LigRMSD 1.0 [63]. The visualization of the ligand-receptor interaction sites was performed with *BIOVIA Discovery Studio 2021*.

## Molecular Dynamics

The ligand-receptor complexes generated with *Autodock Vina* were subjected to molecular dynamics simulations with *YASARA structure v23.12.24* [64]. The simulations start by executing the md_run macro, which included an optimization of the hydrogen bond network to increase the stability of the solute and a pKa prediction to fine-tune the protonation states of the protein residues at the chosen pH of 7.4. NaCl ions were added at a physiological concentration (0.9%), an excess of Na+ or Cl-was added to neutralize the cell. After minimizing using the steepest descent and simulated annealing methods to remove clashes, the simulation was run for 100 nanoseconds using the following force fields: Solute (AMBER14) [52], ligands (GAFF2 [65] and AM1BCC [66]), TIP3P water [67]. A cutoff value of 8 Å was applied for Van der Waals forces (the default value used by AMBER [68]). No cutoff value was applied to electrostatic forces (using the Particle Mesh Ewald algorithm [69]). The equations of motion were integrated every 1.25 fs for bonded interactions and 2.5 fs for non-bonded interactions at a temperature of 310 K and a pressure of 1 atm. After inspecting the root mean square deviation (RMSD) of the solute as a function of simulation time, the first 100 picoseconds were considered as equilibration time and excluded from further analysis. The study of the binding energy using the MM-PBSA method was carried out by executing the md_analyzebindingenergy macro. Directed simulations were performed by executing the md_runsteered macro.

## Cell Culture

Human lung fibroblasts from the primary cell line CCD-8Lu acquired from ATCC (catalog number: CCL-201) were cultured in 100 mm Petri dishes. The cells were maintained in Gibco™ DMEM/F-12 culture medium (catalog number: 11320033). 10% decomplemented fetal bovine serum (FBS) Gibco™ (catalog number: 16000044) and 1% penicillin, amphotericin B and streptomycin from Gibco™ (catalog number: 15240-062) were added to the medium. The cells were incubated at 37°C and 5% CO_2_.

## Induction of Senescence

The induction, standardization, and validation of the replicative senescence (RS) and stress-induced premature senescence (SIPS) models with hydrogen peroxide were reported in previous work [14].

## Drug Preparation

For the biological assays, the following reagents were used: tolvaptan (T7455-5MG, Sigma Aldrich), cryptotanshinone (C5624-5MG, Sigma Aldrich), sotrastaurin (SML3210-5MG, Sigma Aldrich), bicuculline (14340-25MG, Sigma Aldrich), dasatinib (CDS023389-25MG, Sigma Aldrich), quercetin (PHR1488-1G, Sigma Aldrich), dimethyl sulfoxide (DMSO) (D5879-1L, Sigma Aldrich) and 3-(4,5-dimethylthiazol-2-yl)-2,5-diphenyltetrazolium bromide (MTT) (M2128-1G, Invitrogen). All drugs were dissolved in DMSO (100%) at a concentration of 100 mM (stock solution), except cryptotanshinone which was dissolved at 26.99 mM. All dissolved drugs were stored at -20°C. From the stock, dilutions of each drug were prepared in culture medium with 10% FBS without antibiotic at a concentration of 100 μM with 0.1% DMSO. A positive control was prepared using a combination of dasatinib and quercetin (D+Q) of 250 nM D and 50 μM Q, this concentration has been reported in the literature [25]. Medium with 10% FBS without antibiotic and 0.1% DMSO was also prepared. Each solution was filtered with 0.22 μm filters. For the different concentrations at which the senolytic activity was analyzed, serial dilutions were used from the 100 μM solution, such that the only solution with a concentration of 0.1% DMSO (the highest) was the 100 μM of each drug.

## Experimental Validation of the Senolytic Effect Using the MTT Assay

In 96-well plates, 15,000 cells under SIPS (day 6) and RS (24 hours) were seeded. Additionally, 10,000 proliferating cells (passage < 10) were seeded. For cell counting, 10 μl of cell suspension were mixed with 10 μl of trypan blue, from this mixture 10 μl were taken and the TC20™ automatic counter from Biorad (catalog number: 145-0101) was used to count the cells. The cells under RS and proliferating were maintained for 24 hours in incubation at 37°C and 5% CO_2_ before treatment application. After 24 hours the proliferating cells reach a population of approximately 15,000. 200 μl of each drug at each studied concentration, 200 μl of the vehicle (0.1% DMSO) and 200 μl of the positive control (D+Q) were added. The cells were incubated for 48 hours at 37°C and 5% CO_2_, this protocol is common in screenings for senolytics [70]. After the time, the treatments were removed, and the cells were washed 2X with PBS. An MTT solution dissolved in PBS at a final concentration of 0.5 mg/mL and pH 7.4 was prepared and filtered with a 0.22 μm filter. 100 μl of MTT were added to each well and the cells were incubated for 2 hours at 37°C and 5% CO_2_ in complete darkness. After the time, the MTT was removed and 150 μl of DMSO were added, the plates were placed on gentle agitation for 10 minutes and then incubated for 15 minutes at 37°C. The relative optical density was quantified using a Multiskan SkyHigh microplate spectrophotometer (catalog number: A51119500C, Thermo Scientific), the plates were shaken for 10 seconds, and the absorbance was read at 570 nm. For each drug at each concentration, 3 independent experiments were performed in triplicate.

## Analysis of MTT Assay Results

The spectrophotometer readings were processed as follows: The proliferating and senescent controls were normalized to 1, the other groups and treatments were normalized with respect to their control. Next, a Shapiro Wilk normality test and a Levene’s test to assess equality of variances were performed; in both tests, a *p < 0.05* was considered significant. Afterwards, a one-way ANOVA was performed considering a *p < 0.05* as significant, followed by a Tukey’s multiple comparisons test with Benjamini-Hochberg adjustment to reduce the false positive rate, a *p < 0.05* was considered. The data were visualized by comparing the control and senescent groups at different concentrations using bar graphs of the data normalized with respect to the control.

## Caspase-3 Active Immunofluorescence

At the desired timepoint, cells were fixed with 4% paraformaldehyde (PFA) for 15 minutes at room temperature. After this time, PFA was removed, and cells were washed 2X with PBS. PBS was removed, and cells were incubated in PBS with 0.1% Triton X-100 for 45 minutes to permeabilize the cells. After this time, the permeabilization solution was removed and cells were washed 3X with PBS. Next, cells were incubated in PBS with 0.1% Tween and 3% albumin for 90 minutes to block non-specific sites. Subsequently, the blocking solution was removed, and cells were washed 3X with PBS. To measure apoptosis caused by the drugs, the anti-active caspase 3 antibody (ab32042, abcam) was used at a 1:100 dilution. The primary antibody (AB) was diluted in 0.1% PBS-Tween, and each sample was incubated with 50 μl of AB for 24 hours at 4°C with gentle agitation. Subsequently, the AB was removed, and 3X washes with PBS were performed. The secondary ABs were used at a 1:1000 dilution, the following ABs were used: Goat anti-Rabbit IgG (H+L) Cross-Adsorbed Secondary Antibody, Alexa Fluor™ 488 (A11008, Invitrogen). The secondary ABs were diluted in 0.1% PBS-Tween and each sample was incubated with 50 μl of AB for 1 hour at room temperature in complete darkness and without agitation. After this, the secondary AB was removed and 3X washes with PBS were performed. The mounting solution was prepared using 500 μl of distilled water and 500 μl of glycerol to which 1 μl of 4′,6-diamidino-2-phenylindole (DAPI) (D1306, Invitrogen) was added, this process was carried out in complete darkness. The coverslips with the cells were mounted onto 50 μl of mounting solution on glass slides for 25 minutes and subsequently sealed with varnish.

## Quantification and Statistical Analysis of Fluorescence

Using the Zeiss AXIOVERT 40 CFL microscope and the *ZEN 2.3 (Blue edition)* software, photographs of 3 different slides per group were taken. The 60x magnification was used, capturing approximately the same cell density for both controls and experimental groups using the same exposure. For each slide, 3 photos of different sections were taken. Fluorescence quantification was performed using ImageJ software, briefly, the algorithm does the following: 1) converts images to grayscale; 2) removes background noise using a background photo from each group; 3) quantifies total pixel intensity and subtracts background; 4) performs a normality test; 5) performs an independent student’s t-test (*p < 0.05* was considered significant); and 6) generates a graph comparing the two groups.

## Author Contributions

S.O.: Conceptualization, computational analyses, experimental validation, figure design, and article writing. M.K., J.P.V, N.E.L.D: Conceptualization, project direction, and advisement, funding and material resources, writing, editing, review, and article approval. F.C.B.: Advisement on computational analyses, writing and review article. All authors approved the final version of the manuscript.

## Funding

This work was supported by Consejo Nacional de Humanidades, Ciencias y Tecnologías (CONAHCYT) grant FORDECYT-PRONACES/263957/2020. S.O. is a CONAHCyT scholarship holder.

## Conflict of Interest Statement

The authors of this study are currently in the process of registering an international patent to protect the senolytic activity discovered for the molecules reported in this work.

## Acknowledgments

We thank CONAHCyT for the doctoral scholarship awarded to Samael Olascoaga CVU: 869300.

We thank the Laboratorio de Supercómputo y Visualización en Paralelo (LSVP) of the Universidad Autónoma Metropolitana Campus Iztapalapa for the computing time granted on the ’Yoltla’ supercomputer.

## Data availability

All data generated or analyzed during this study are included in this published article in the supplementary files.

## Code availability

The code used to support the analysis of this study can be found at https://github.com/Olascoaga/Senolytic-Repurposing

