## Supplementary material for "Discovery and Repurposing of Multi-Target Senolytics through Structure-Based Virtual Screening": S1

### Supplementary Tables and Figures (S1)

Database of senolytics reported in literature

| Name | Weight | Structure | Ref |
| --- | --- | --- | --- |
| 2,3-Dimethoxy-1,4-naphthoquinone | 188.18 | 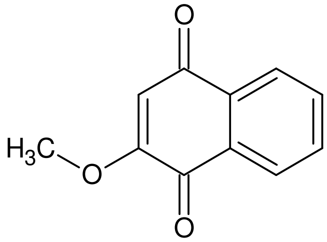 | 1 |
| A1155463 | 669.80 | 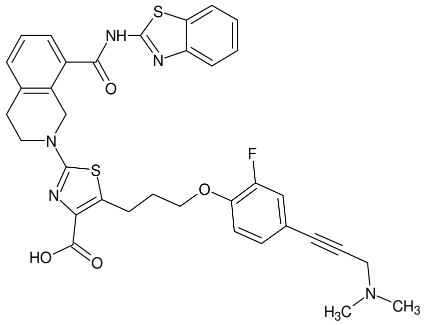 | 1 |
| A1331852 | 658.80 | 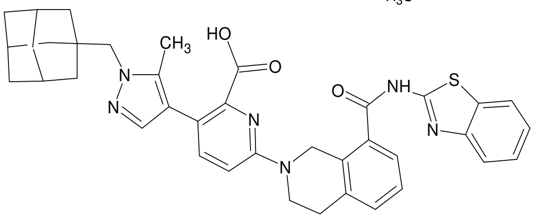 | 1 |
| ABT-737 | 813.40 | 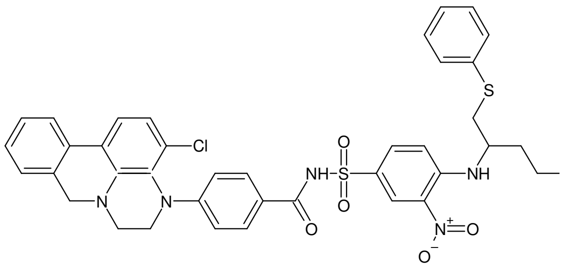 | 1 |
| Alvespimycin | 616.70 | 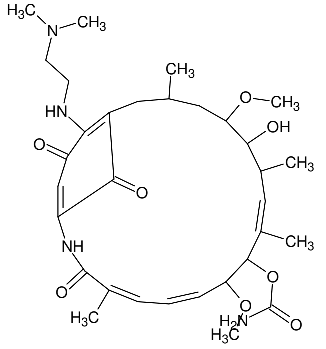 | 1 |

| | | | |
|---|---|---|---|
| Analoge 49 | 311.33 | 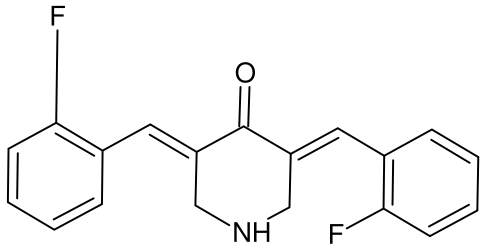 | 1 |
| Antipyrine | 188.23 | 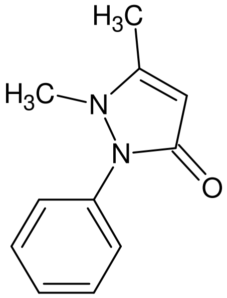 | 1 |
| Astemizole | 458.60 | 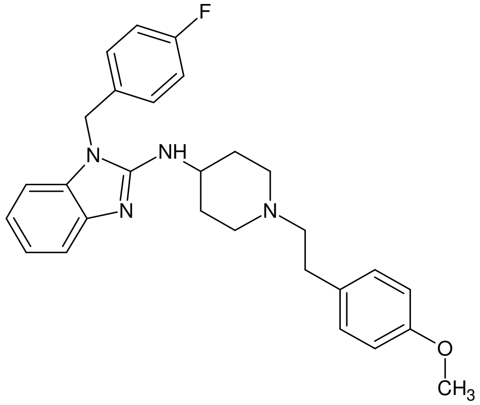 | 1 |
| Azacyclonol | 267.37 | 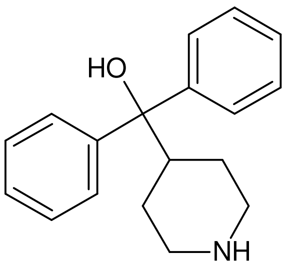 | 2 |
| Azithromycin | 749.00 | 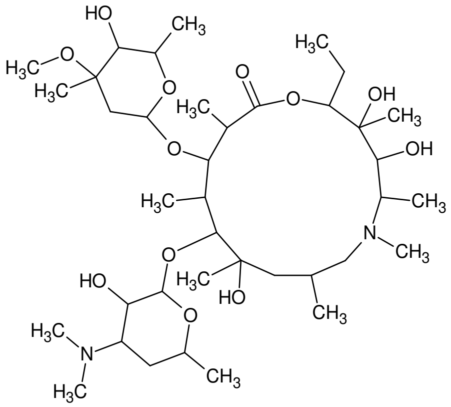 | 2 |

| | | | |
|---|---|---|---|
| Benzydamine hydrochloride | 309.41 | 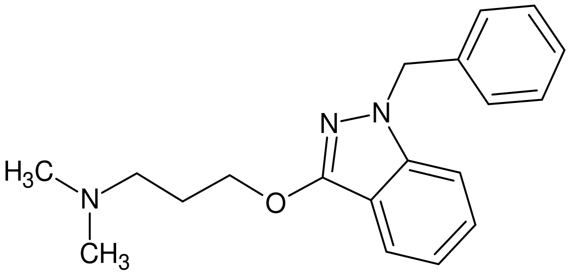 | 2 |
| Berberine | 336.37 | 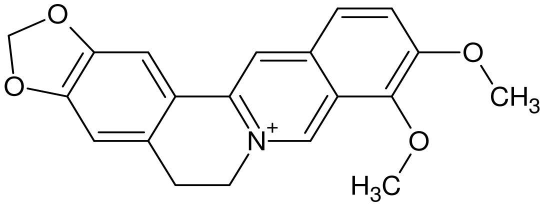 | 2 |
| BIIB021 | 318.77 | 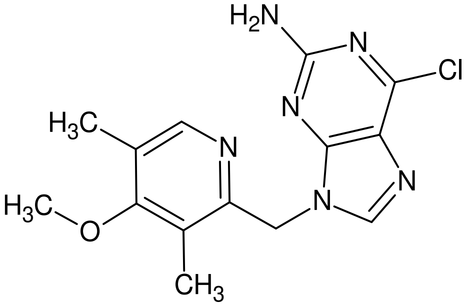 | 2 |
| Bupivacaine hydrochloride | 288.43 | 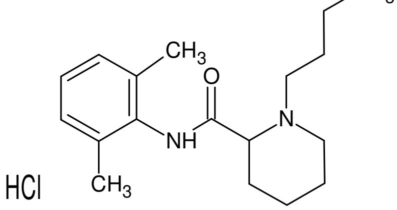 | 2 |
| Cantharidin | 196.20 | 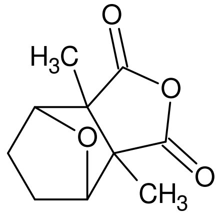 | 2 |
| Ciproheptadine hydrochloride | 287.41 | 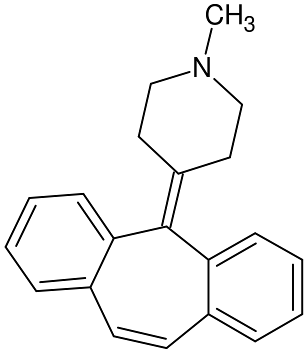 | 2 |
| Curcumin | 368.38 | 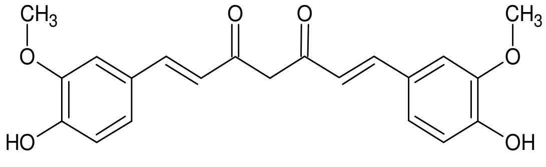 | 2 |

| | | | |
|---|---|---|---|
| Dasatinib | 488.01 | 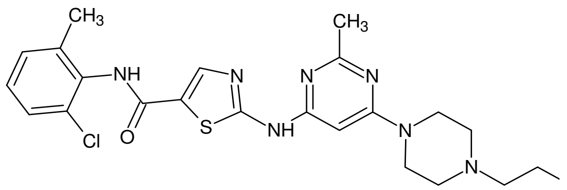 | 2 |
| Digitoxigenin | 374.52 | 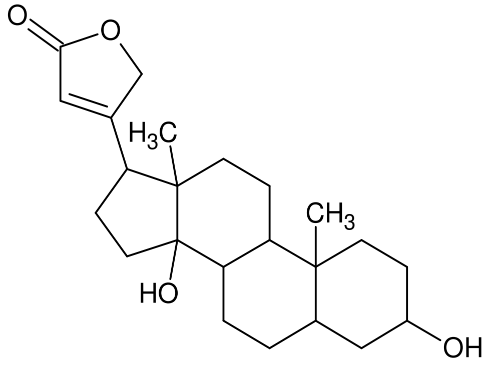 | 2 |
| Digoxin | 780.90 | 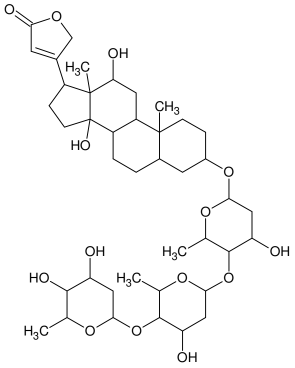 | 2 |
| Diphenyleneiodonium | 314.55 | 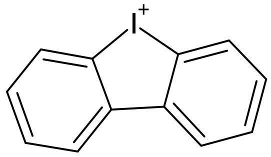 | 3 |
| Eltrombopag | 442.47 | 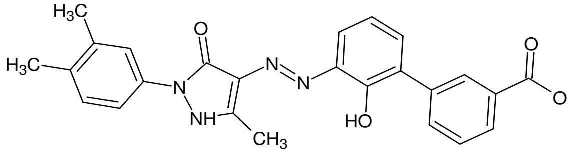 | 3 |
| Fenofibrate | 360.84 | 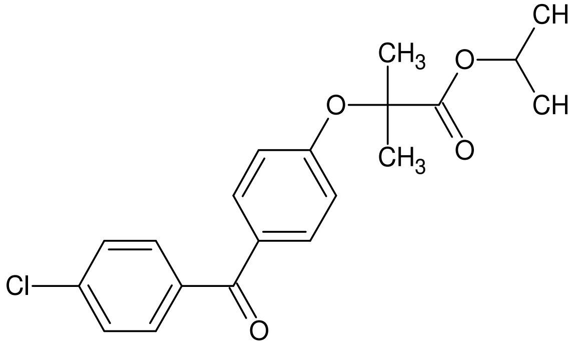 | 4 |

| | | | |
|---|---|---|---|
| Fisetin | 286.24 | 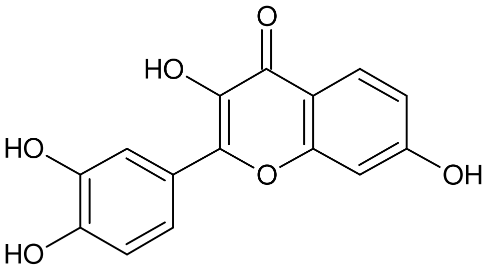 | 4 |
| Flutamide | 276.21 | 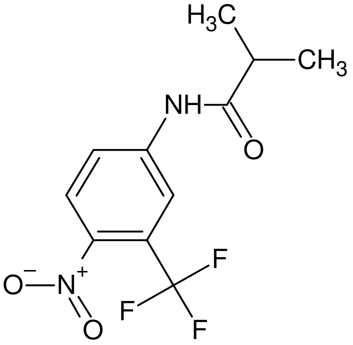 | 4 |
| Ganetespib | 364.40 | 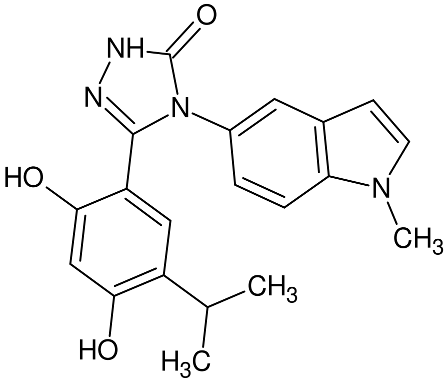 | 4 |
| Geldanamycin | 560.60 | 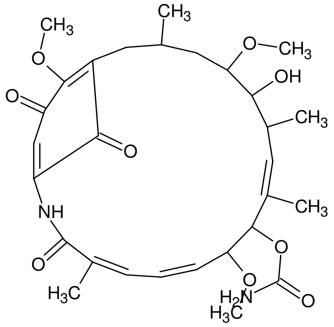 | 4 |
| Gemcitabine | 263.20 | 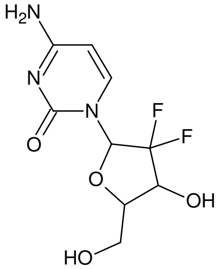 | 5 |
| Genistein | 270.24 | 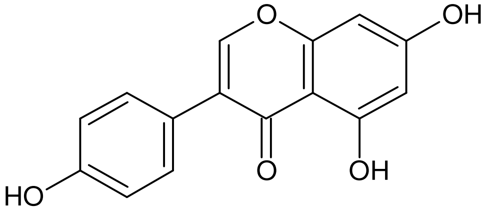 | 6 |

| | | | |
|---|---|---|---|
| Guanethidine | 198.31 | 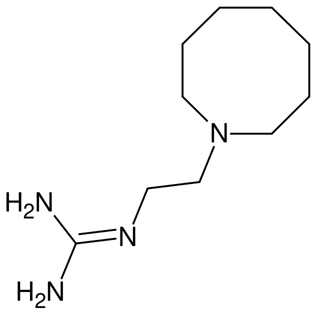 | 6 |
| Idarubicin | 497.50 | 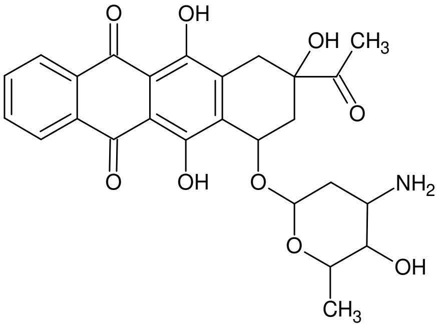 | 7 |
| Lomefloxacin | 387.80 | 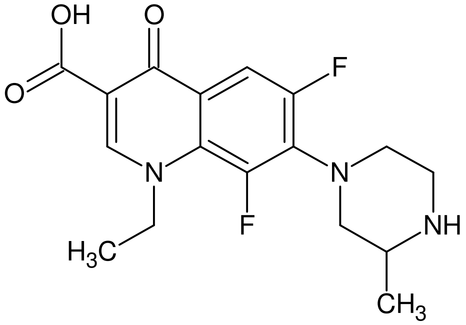 | 8 |
| LY-367,265 | 452.55 | 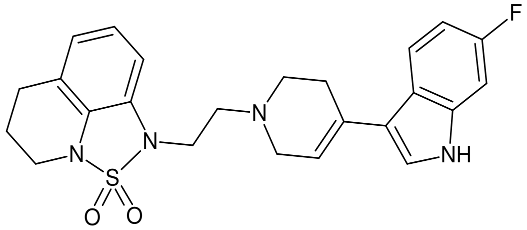 | 9 |
| Minoxidil | 289.32 | 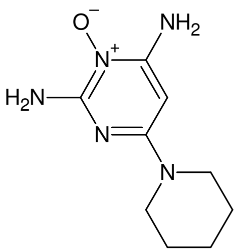 | 9 |
| MK2206 | 407.50 | 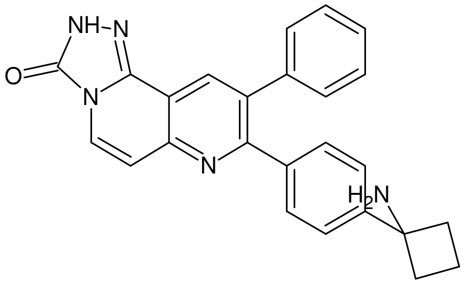 | 10 |

| | | | |
|---|---|---|---|
| Navitoclax | 974.61 | 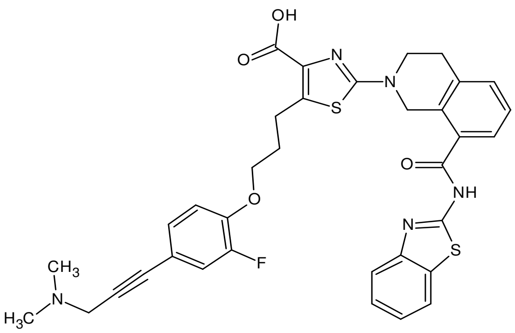 | 10 |
| Nitazoxanide | 307.29 | 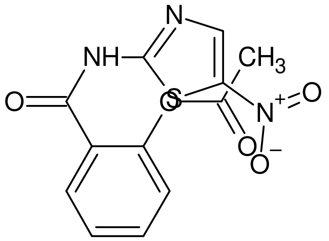 | 11 |
| Nitrofurazone | 198.14 | 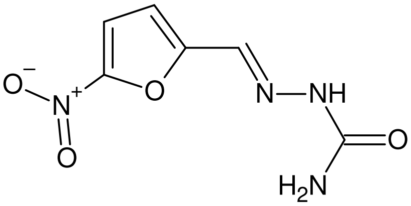 | 11 |
| Ouabain | 584.70 | 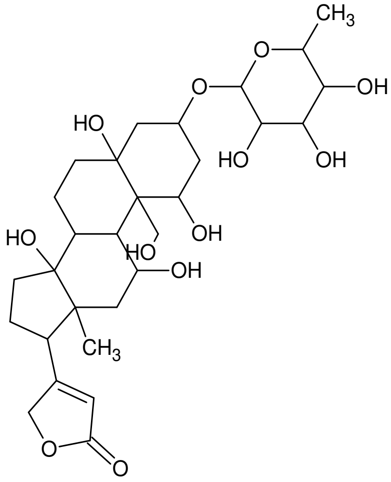 | 11 |
| O-VANILLIN | 152.15 | 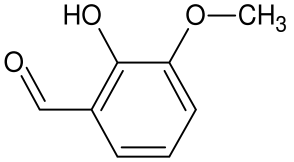 | 11 |
| Panobinostat | 349.43 | 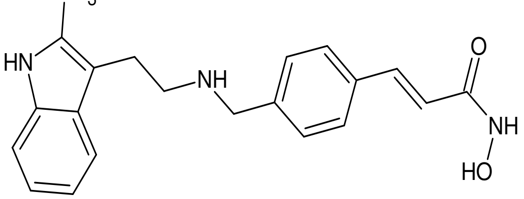 | 11 |

| | | | |
|---|---|---|---|
| Pherphenazine | 403.97 | 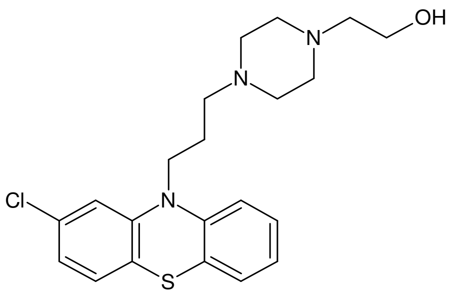 | 11 |
| Phloretin | 274.27 | 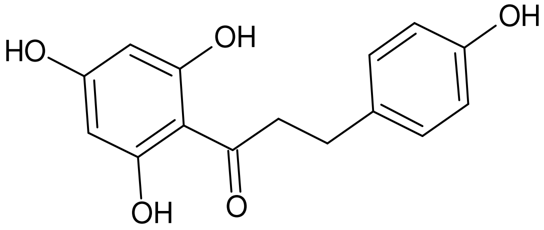 | 11 |
| Piperlongumine | 317.34 | 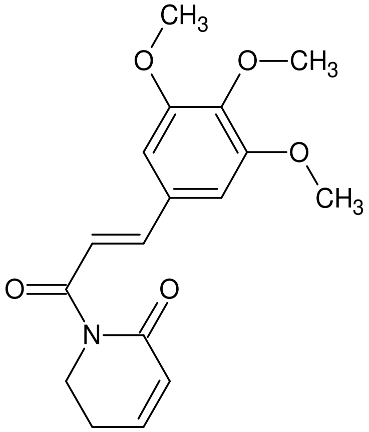 | 12 |
| Proscillaridin A | 530.60 | 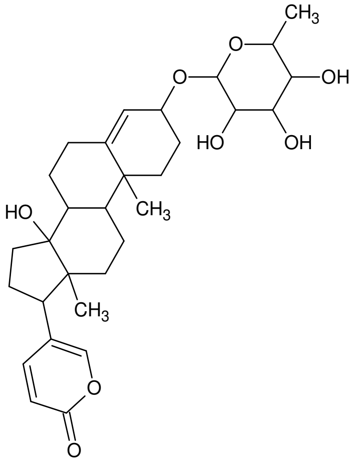 | 13 |
| Quercetin | 302.24 | 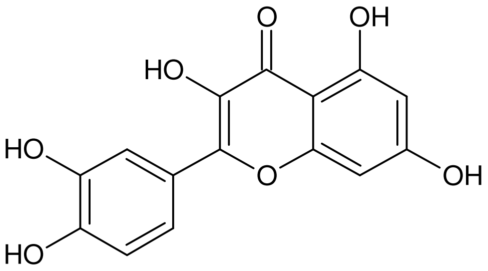 | 14 |

| | | | |
|---|---|---|---|
| R406 | 470.5g | 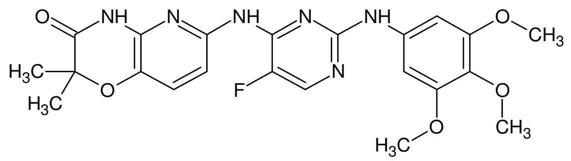 | 15 |
| Radicicol | 364.78 | 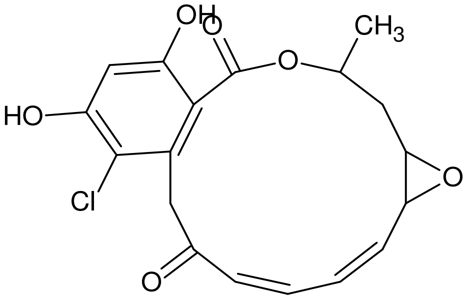 | 16 |
| Rottlerin | 516.54 | 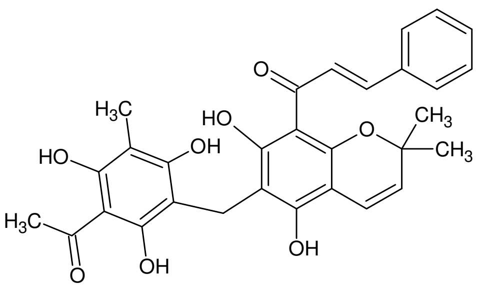 | 16 |
| Roxithromycin | 837.00 | 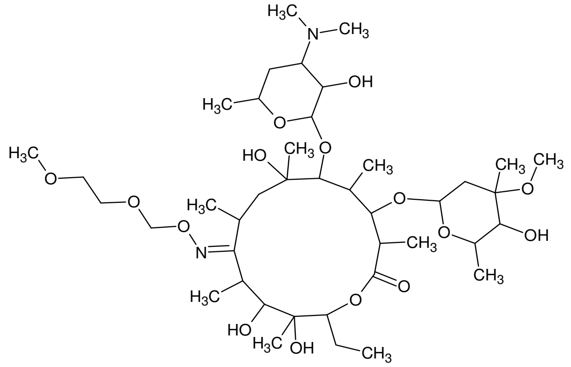 | 17 |
| Tanespimycin | 585.70 | 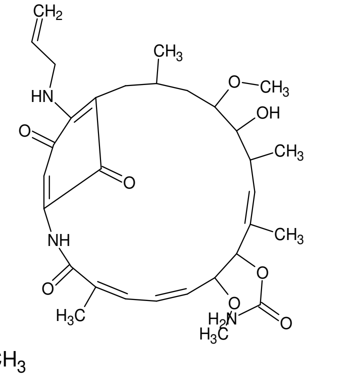 | 17 |
| Troglitazone | 441.55 | 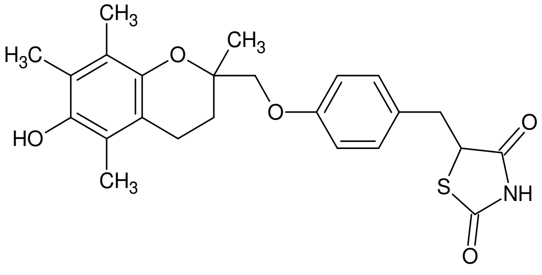 | 17 |

| | | | |
|---|---|---|---|
| Tyrphostin | 315.76 | 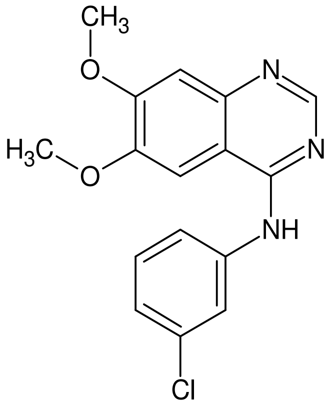 | 18 |
| Vorinostat | 264.32 | 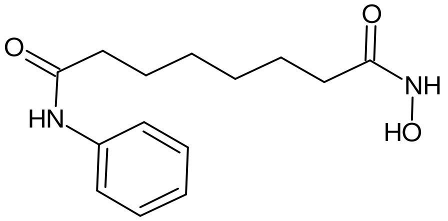 | 19 |

| | | | |
|---|---|---|---|
| Alectinib | 482.60 | 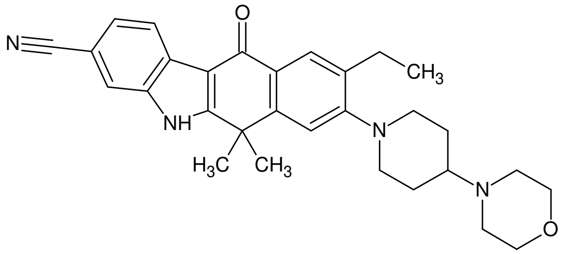 | 20 |
| Luminespib | 465.50 | 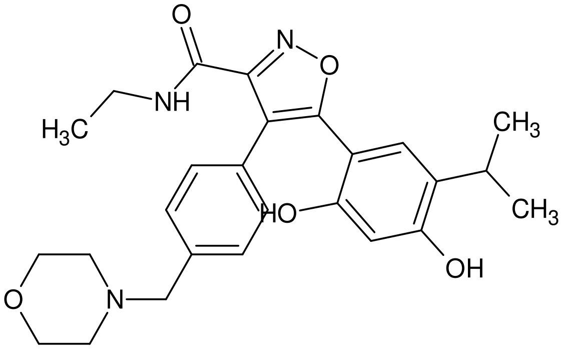 | 21 |
| Onalespib | 409.50 | 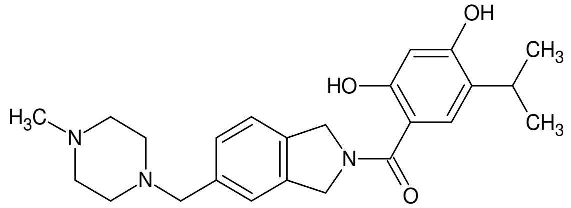 | 22 |
| Cycloheximide | 281.35 | 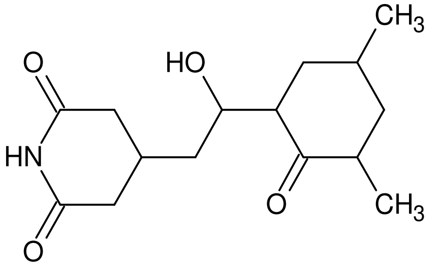 | 23 |
| WEHI-539 | 583.70 | 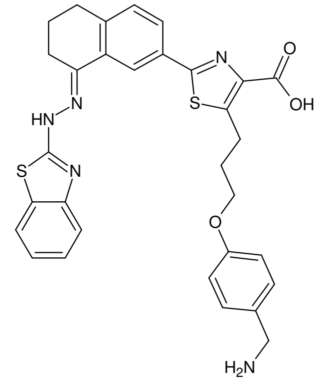 | 24 |
| Epigallocatechin gallate | 458.40 | 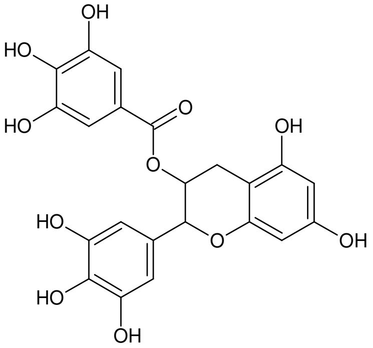 | 25 |

| | | | |
|---|---|---|---|
| Gama-Tocotrienol | 410.60 | 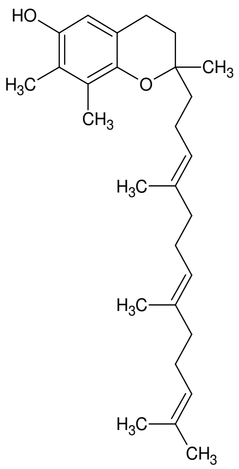 | 26 |
| Ginsenoside Rg1 | 801.00 | 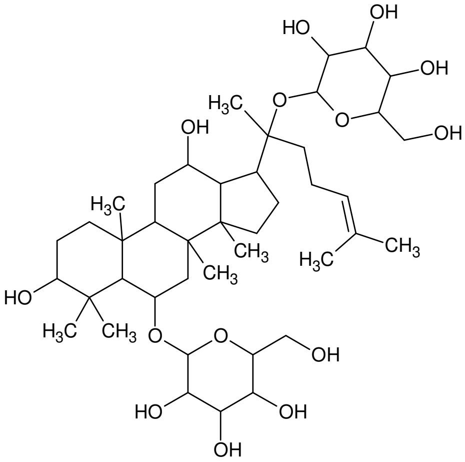 | 27 |
| Spermidine | 145.25 | 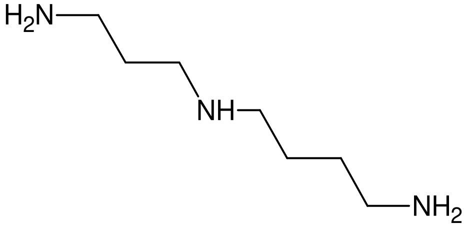 | 28 |
| EF24 | | 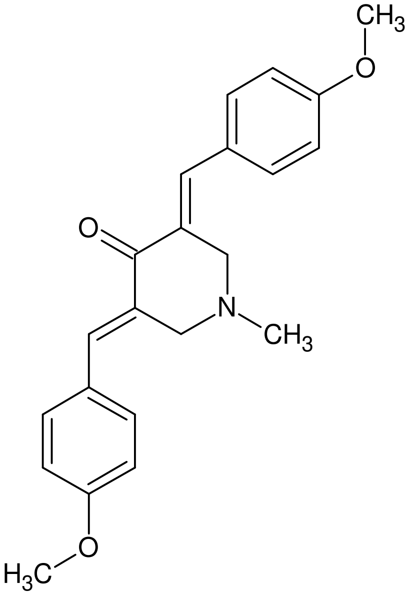 | 29 |

| | | | |
|---|---|---|---|
| PU-H71 | 512.40 | 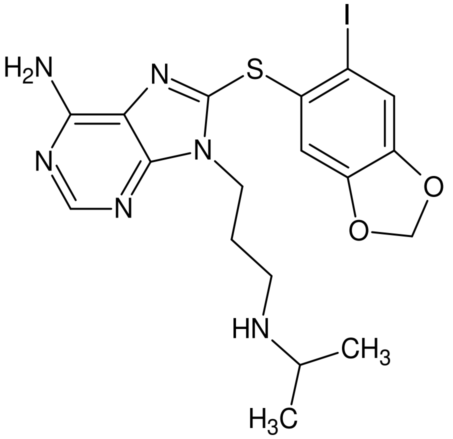 | 30 |
| Luteolin | 286.24 | 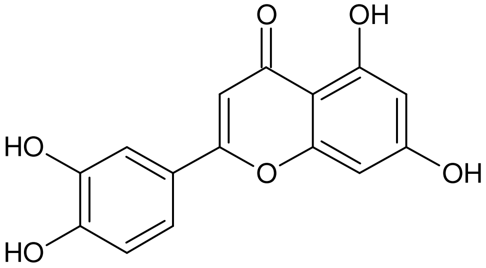 | 30 |
| Hydroxytyrosol | 154.16 | 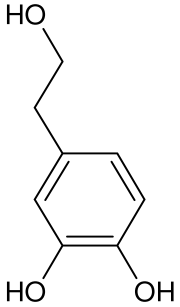 | 31 |
| Hyperoside | 464.40 | 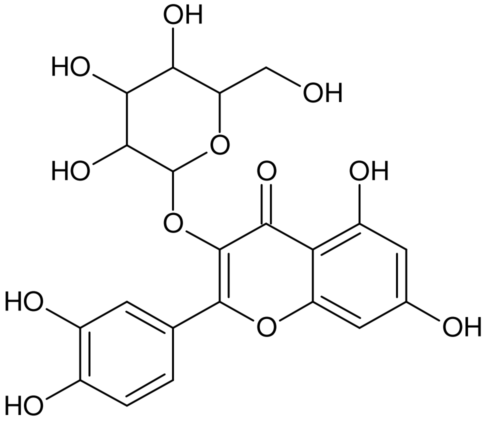 | 32 |
| Alliin | 177.22 | 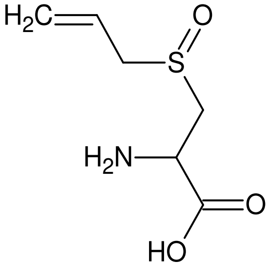 | 33 |

| | | | |
|---|---|---|---|
| Allicin | 162.30 | 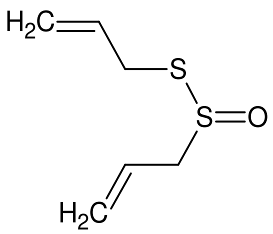 | 33 |

**Table S1.** Database of 70 senolytics reported in literature, with name, molecular weight, structure and reference.

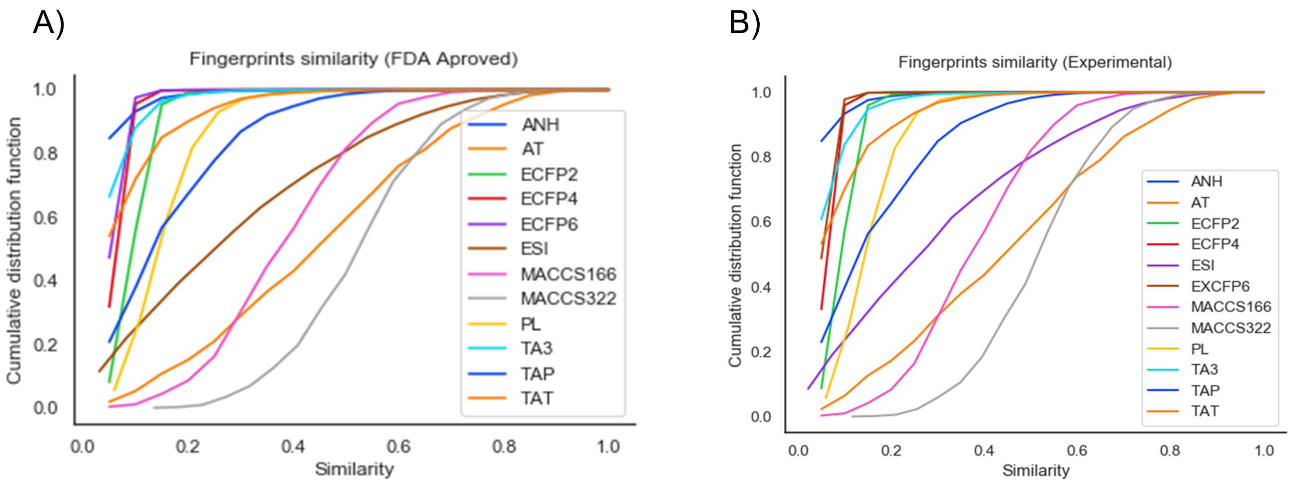


**Figure S1.** Cumulative distribution curves of 12 molecular fingerprints (FP). Path Length (PL), MACCSKeys 166 and 322 bits (MACCS), Atom Neighborhoods (ANH), AtomTypes (AT), EStateIndices (ESI), Extended Connectivity radius 2, 4 and 6 (ECFP2, 4, 6), Topological Atom Pairs (TAP), Topological Atom Torsions (TAT) and Topological Atom Triplets (TA3) evaluated in A) FDA-approved drug database and B) experimental phase drugs.

| *FP* | Min | Max | Mean | Median | STD |
|---|---|---|---|---|---|
| *ANH* | 0 | 1 | 0.02 | 0 | 0.05 |
| *AT* | 0 | 1 | 0.44 | 0.44 | 0.21 |
| *ECFP2* | 0 | 1 | 0.09 | 0.09 | 0.04 |
| *ECFP4* | 0 | 1 | 0.06 | 0.05 | 0.03 |
| *ECFP6* | 0 | 1 | 0.05 | 0.05 | 0.02 |
| *ESI* | -0.02 | 1 | 0.29 | 0.25 | 0.22 |
| *MACCS166* | 0 | 1 | 0.38 | 0.37 | 0.13 |
| *MACCS322* | 0.09 | 1 | 0.52 | 0.52 | 0.13 |
| *PL* | 0.01 | 1 | 0.15 | 0.14 | 0.07 |
| *TA3* | 0 | 1 | 0.04 | 0.03 | 0.05 |
| *TAP* | 0 | 1 | 0.16 | 0.13 | 0.12 |
| *TAT* | 0 | 1 | 0.07 | 0.04 | 0.09 |

**Table S2.** Descriptive parameters for each FP of the FDA-approved drug database

| *FP* | Min | Max | Mean | Median | STD |
|---|---|---|---|---|---|
| *ANH* | 0 | 1 | 0.02 | 0 | 0.05 |
| *AT* | 0 | 1 | 0.44 | 0.44 | 0.22 |
| *ECFP2* | 0 | 1 | 0.09 | 0.09 | 0.04 |
| *ECFP4* | 0 | 1 | 0.06 | 0.05 | 0.02 |
| *ESI* | -0.03 | 1 | 0.30 | 0.26 | 0.22 |
| *EXCFP6* | 0 | 1 | 0.05 | 0.05 | 0.02 |
| *MACCS166* | 0 | 1 | 0.37 | 0.37 | 0.13 |
| *MACCS322* | 0.07 | 1 | 0.51 | 0.51 | 0.13 |
| *PL* | 0.01 | 1 | 0.15 | 0.14 | 0.07 |
| *TA3* | 0 | 1 | 0.05 | 0.03 | 0.06 |
| *TAP* | 0 | 1 | 0.16 | 0.13 | 0.13 |
| *TAT* | 0 | 1 | 0.07 | 0.04 | 0.09 |

**Table S3.** Descriptive parameters for each FP of the experimental phase drug database

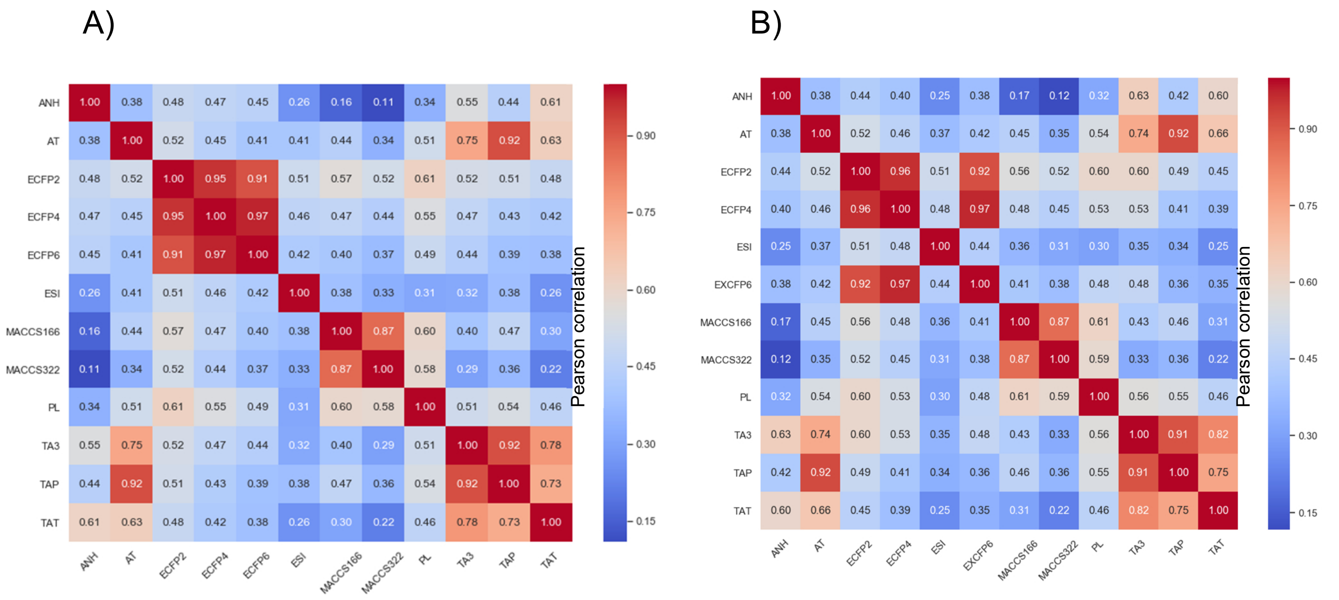


**Figure S2.** Pearson correlation analysis of similarity matrices calculated with the Tanimoto index of 12 different FPs for A) FDA-approved drugs and B) Experimental phase drugs.

| Drug | Structure | Targets | DB | Original Prescription |
|---|---|---|---|---|
| Sotrastaurin | 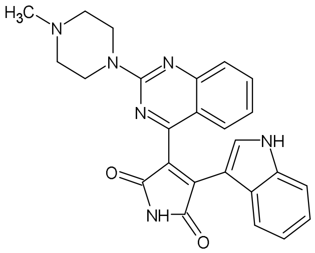 | PDGFB, SERPINE1, PIK3CD, EFNB1 | EXP | Kidney and liver transplantation<br><br>Moderate Psoriasis<br><br>Ulcerative Colitis |
| Drospirenone | 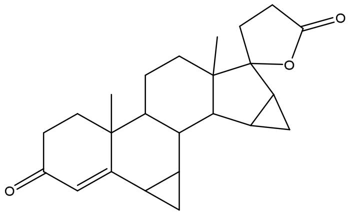 | SERPINE1, EFNB1, PIK3CD | FDA | Anticonceptive<br><br>Polycystic Ovarian Syndrome |
| Peruvoside | 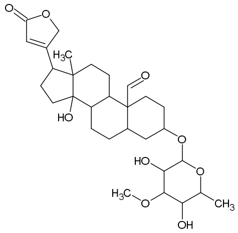 | PDGFB, PIK3CD, EFNB1 | EXP | N/A |
| Tolvaptan | 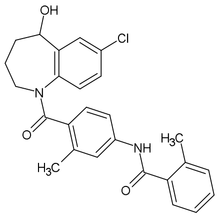 | PDGFB, PIK3CD, EFNB1 | FDA | Hyponatremia<br><br>Heart Failure |
| Amrubicin | 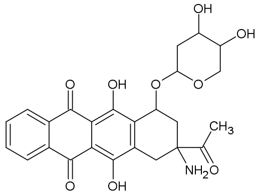 | PDGFB, SERPINE1, EFNB1 | EXP | Lung cancer |
| Cryptotanshinone | 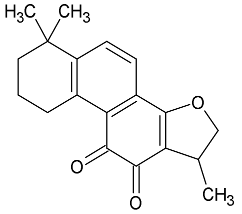 | SERPINE1, PIK3CD | EXP | N/A |
| Dicoumarol | 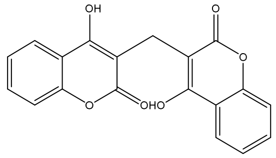 | PDGFB, SERPINE1 | FDA | Anticoagulant |

| | | | | |
|---|---|---|---|---|
| Naldemedine | 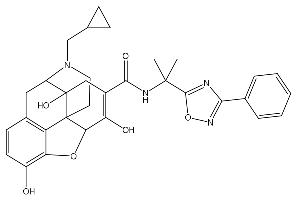 | PIK3CD, EFNB1 | FDA | Opioid Induced constipation |
| Bicuculline | 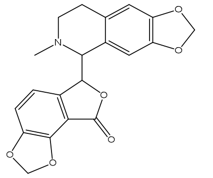 | SERPINE1, EFNB1 | EXP | N/A |
| Ciclesonide | 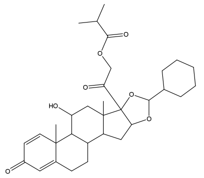 | PDGFB, EFNB1 | FDA | Asthma<br><br>Allergic Rhinitis |
| Azatadine | 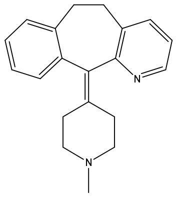 | SERPINE1, EFNB1 | FDA | Antihistaminic |

**Table S4**. Top 11 drugs with the highest multi-target potential.

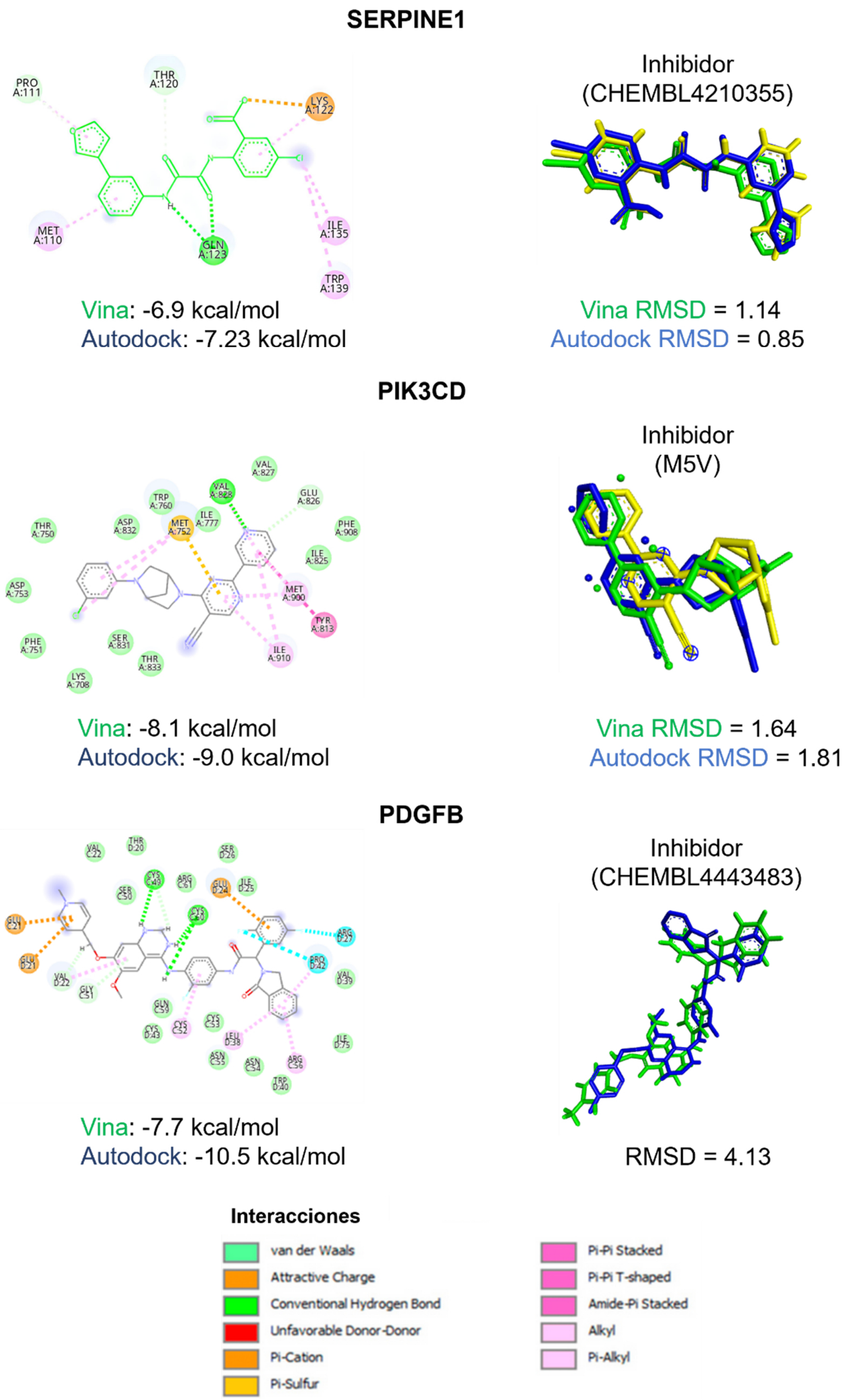


**Figure S3**. Validation of the molecular docking protocol. "Re-docking" of co-crystallized inhibitors reported in the literature was performed, and the RMSD of the co-crystallized structure (yellow) and the best energetic position generated with Vina (green) and Autodock4 (blue) was quantified.
